# Sequence and structural analysis of adaptors of Toll-like receptor 4 sheds light on the evolutionary trajectory and functional emergence

**DOI:** 10.1101/2024.07.30.605793

**Authors:** Shailya Verma, Ramanathan Sowdhamini

## Abstract

Type 4 Toll-like receptors 4 (TLR4) recognise lipopolysaccharides (LPS) from bacteria as their conventional ligands and undergo downstream signalling to produce cytokines. They mediate the signalling either by the TIRAP-MyD88 complex or by the TRAM-TRIF complex. The MyD88 pathway is common to all other TLRs, whereas the TRAM-TRIF complex is largely exclusive to TLR4. We studied the TIR domain of TRAM and TRIF homologue proteins, that are crucial for downstream signalling. From our previous work on pan-genome-wide survey, we find *Callorhincus milli* to be the ancestral organism with both TRAM and TRIF proteins. To gain a deeper insight about the functioning of these proteins and comparison with the adaptor proteins in *Homo sapiens*, we performed TRAM and TRIF dimer docking to model the TRAM-TRIF complex of representative organisms across various taxa. These provide us insights to ascertain a possible interaction surface, calculate the energetics, electrostatic potential, and then employ Normal Mode Analysis (NMA) to examine fluctuating, interacting and specific residue clusters which can be important for the protein functioning in both organisms. We also performed molecular dynamics simulations of these complexes and cross validated the functionally important residues using network parameters. While the critical residues of TIRAP, TRIF, and MyD88 were preserved, we found that the important residues of TRAM signalling were not conserved in *Callorhincus milli*. This suggests the presence of functional TIRAP-MyD88 mediated TLR4 signalling and TRIF mediated TLR3 signalling in the ancestral species. The overall biological function of this signalling domain appears to be gradually acquired through the orchestration of several motifs through evolutionary scale.

## 2. Introduction

Toll like receptors are key pattern recognition receptors responsible for identification of various pathogen associated molecular patterns (PAMPs) and danger associated molecular patterns (DAMPs). They are part of the innate immune system and protect us from various pathogens. The TLRs are localised on the plasma membranes or on the endosomes. They have an extracellular domain (ECD), that recognizes or interacts with ligand, a transmembrane and an intracellular Toll/interleukin 1 receptor (TIR) domain. TLR4 identifies Lipopolysaccharide (LPS) as its ligand and undergoes two routes of downstream signalling. The conventional path of downstream signalling is by the interaction of TIR adaptor protein (TIRAP) and Myeloid Differentiation Primary Response Protein 88 (MyD88) mediated, that is common for all the plasma membrane localized TLRs. Whereas TLR4 and TLR2 also mediate by the TIR domain-containing adapter molecule 2 (TICAM2/TRAM) and TIR domain-containing adapter molecule 1 (TICAM1/TRIF) to produce type I Interferon IFNs [1] [2]. The Cluster of differentiation 14 (CD14) protein localizes in the membrane and binds to the LPS to bring it to the TLR4 which further recognizes LPS with help from Myeloid differentiation factor-2 (MD2) protein and undergo dimerization. The MyD88 pathway and TRAM mediated pathway are competitive and mutually exclusive to each other and the latter gets activated when the complex internalizes into endosome [3]. Recently, we had reported the evolution of TRAM and TRIF protein across various taxa from tree of life [4]. The conservation pattern of several key residues and motif patterns among orthologs suggest a potential pathway in non-mammals. These are the AEDD site (A85, E86, D87, D88), which is important for upstream interaction of TRAM TIR with TLR4 TIR, BB loop residues that is crucial for TRAM dimer formation, TS site (T155, S156), important for TRAM TIR downstream interaction with TRIF TIR domain and Y167 phosphorylation site [5][6]. Mutations at two residues in the BB loop, P116 and C117, leads to abrogation of downstream signalling [7][8]. Human TRAM has a Myristoylation motif that is responsible for the localization of the protein to the plasma membrane [9]. Also, TRAF 6 binding motif on TRAM ensures activation of response by TLR4. The schematic of TLR-4 pathway with key residues is shown in **Figure 1**.

**Figure 1:**
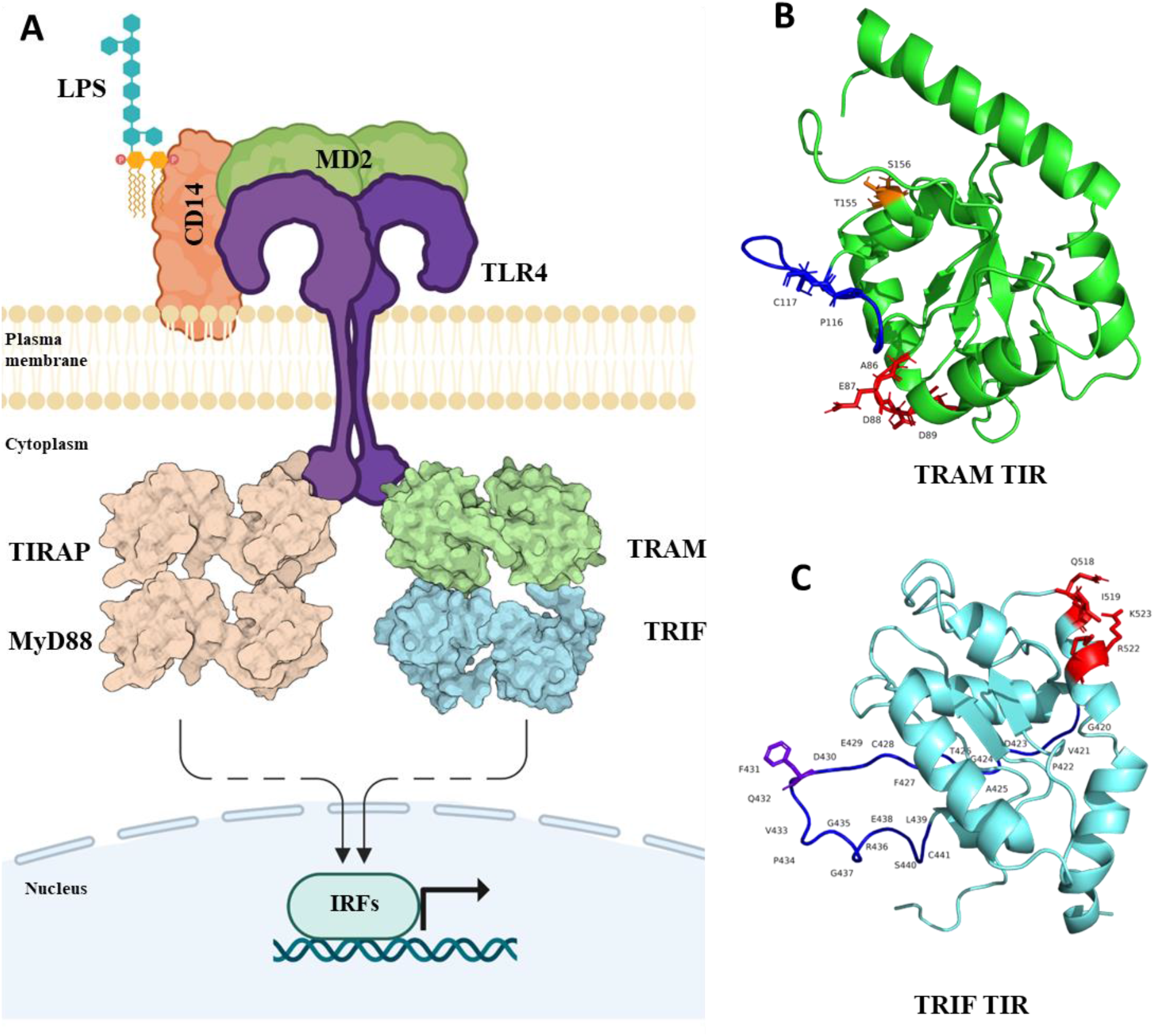
A) Schematic of Toll-like receptor 4 signalling with the adaptor molecules, TIRAP, MyD88, and TRAM, TRIF mediated pathways. The dashed arrow represents multiple other mediators in between that ultimately leads to IRFs (IRF3, IRF7) productions and expression of type I interferon (IFN α/β) genes. B) Structure of TRAM TIR domain with highlighted key residues (AEDD site: red, BB loop: blue, P116 and C117: blue, TS site: orange). C) Structure of TRIF TIR with highlighted key residues (QI and RK site: red, BB loop: blue, F431: purple)

On the other hand, human TRIF protein has three domains, N-terminal domain (NTD), TIR and RIP homotypic interaction motif (RHIM) domains. NTD domain is majorly responsible for nuclear factor kappa B (NF-κB) and interferon regulatory factor 3 (IRF3) production. And the RHIM domain is totally responsible for the TRIF induced apoptosis [10]. TRIF also contains pLxIS motif that contains the phosphorylation site used for IRF3 activation, it also contains TRAF6 binding motif that activates NF-κB production and RHIM motif in the RHIM domain [11]. TRIF TIR has QI (Q518, I519) and RK (R522, K523) site important for interaction with TRAM TIR, It also has F431 that is identified as a key residue for interaction of TRIF TIR with its TRIF-NTD in an autoinhibitory state [5][12].

We have studied the conservation of the key residues in the functional TIR domain of both the proteins, that is crucial for dimer formation and downstream signalling. We have modelled these TRAM TRIF complexes assuming both trimeric and tetrameric orientations. We further used the best model to hypothesize the interaction pattern in case of representative organism across various taxa. We have also used Normal mode analysis and molecular dynamics simulation to understand the complexes better to examine if the primitive orthologs have the functional capability for signal transduction.

## 3. Results

### 3.1 Residue conservation in TRAM and TRIF orthologs

In our analysis we have thoroughly compared and selected few orthologs sequences to compare detailed structural interaction with reference to *Homo sapiens*. We have picked representative organism across various taxonomical orders ranging from the common ancestor of both adaptors in Chondrichthyes, to representatives from Amphibians, Cryptodira, Crocodylia, Bifurcata, Aves and Mammalia. These orthologs were carefully chosen to have well defined domain annotation of adaptors. But before focussing on these particular cases, we did a comparison across all the available sequence of TRAM and TRIF adaptors. As the NMR structure of the TIR domain was available for both TRIF and TRAM protein in PDB (2M1X, 2M1W), we were interested to investigate if the conserved residues amongst the orthologs have any structural significance [13]. Evolutionary Trace method was used to rank functional and structurally important residues among orthologs [14]. A higher score represents sequence position variation among distant orthologs and a lower score represents variation among close orthologs. This prediction was also extended to ConSurf analysis to substantiate the degree of conservation of residues and map it onto the protein structure [15]. Co-evolving residues were also predicted using VisualCMAT [16]. These results are shown by mapping onto the respective secondary structure from PDBsum in **Figure 2** [17].

**Figure 2:**
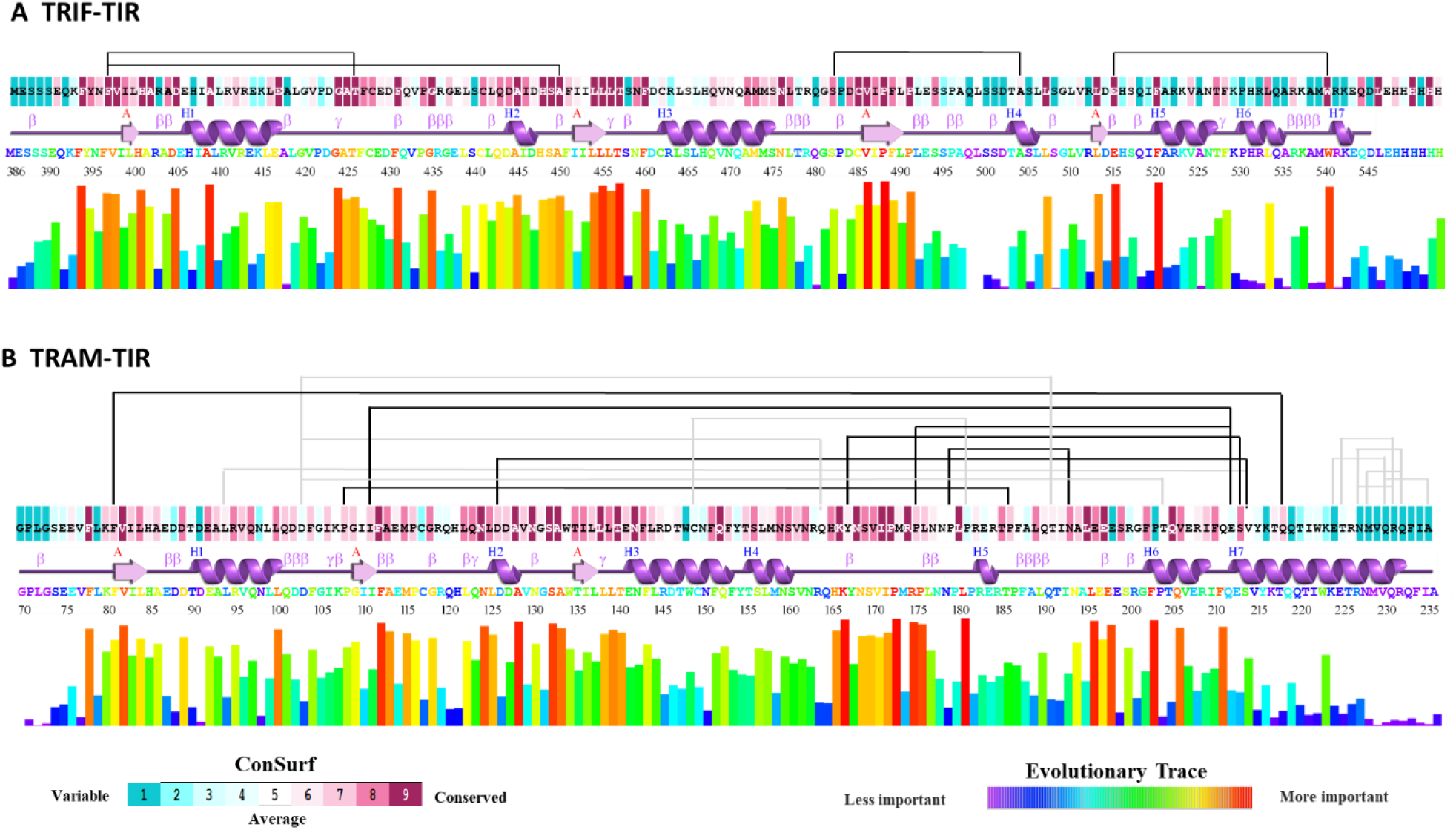
First colour plot shows the ConSurf result for degree of conservation on a scale of 1 to 9 amongst the orthologs. Below is the secondary structure of the TIR domain of the corresponding protein. Then the coloured representation of amino acid sequence and the histogram shows the result from Evolutionary trace method. Above all these the connecting lines between the amino acid shows the co-evolving pairs as predicted from VisualCMAT. **A** TRIF-TIR data being depicted on PDB:2M1X, **B** TRAM-TIR data depicted on PDB: 2M1W.

From these results, we were able to see the evolutionary trace analysis goes in parallel with the ConSurf conserved residue analysis. Also, the result from visualCMAT shows many pairs that have sequences from both highly conserved and variable regions. There were four pairs predicted for TRIF-TIR, and all the residue involved was highly conserved among orthologs. These pairs include F397-T426, F397-A450, S482-A504, and E515-W540. Interestingly, F397 was found to coevolve with T246 as well as A450.

Apart from these, eighteen pairs were predicted to co-evolve amongst TRAM orthologs. From that, we have highlighted the highly conserved residues of co-evolving pairs. These were between F81-218Q, 108P-186P, 111I-212E, 126D-214V, 167Y-213S, 175P-212E and 179P-193N. Amongst these, 212E was seen to coevolve with 111I, as well as 175P. Besides that, 108P, 111I, 126D were near the BB loop regions, so they may have some significant changes in protein function. Also, residues 167Y which is important for response to LPS by phosphorylation was found to be co-evolving with 213S [18]. Among these pairs 212E, 213S, and 214V all seem to co-evolve, this region may also be of some functional relation. Apart from this, the available mutagenesis studies show the significance of P116H, C117H, Y154F, and Y167F [18][7][8]. It will be interesting to investigate the role of these evolving residues and check for their functional role or if it interrupts protein stability.

Further, the list of the co-evolving residues along with other parameters is provided in Additional File 2, Table S1. Few other combinations of co-evolving residues and their presence in the orthologs at those positions respectively. Also, the frequency of occurrence of such varied amino acids is shown in the table. For those combinations that had frequency =>1%, we performed virtual mutations at respective positions on *Homo sapiens* TIR structure and calculated the free energy change (ΔΔG kcal/mol = ΔGmut-ΔGwt) using FoldX 5.0 [19].

In TRIF-TIR domain, apart from F397-A450, other pairs were mostly coevolving either to neutral or toward stabilising end. In TRAM-TIR domain, most of the mutations lead to destabilising or neutral except for D126-T204.While analysing the homologs sequences for coevolving residue pair we found corresponding Phenlyalanine at 81^st^ position of TRAM and 397^th^ of TRIF is part of coevolving pair. This residue is highly conserved and is found to be destabilized when mutated to other residue as observed in orthologs sequences. Thereby, this residue seems to play an important role in the TIR functioning of both adaptors. Apart from this Y167 from TRAM which is known to be important for phosphorylation function coevolves with S213 and leads to either destabilising or neutral mutations in orthologs. The details of the free energy change due to coevolving mutation in TRAM and TRIF is mentioned in Additional File 1, Figure S1.

### 3.2 Important motifs and residue conservation in chosen representative organism

While focussing on the various taxa representatives, it becomes imperative to observe the key residue patterns, major motif and domain conservation to ascertain the protein function. We thereby looked at the multiple sequence alignment of representative organism and mapped the key residues, motifs and other patterns of TRAM (AEDD, PC, TS, Y site) and TRIF (QI and RK site) protein [5]. We also looked at domain architecture and key Myristoylation, TRAF6 binding site motif of TRAM protein and pLxIS motif, TRAF6 binding motifs of TRIF protein. The **Figure 3** below shows the conservation pattern of the key residues in the representative orthologs. We also observed the conservation pattern in MyD88 and TIRAP protein to validate the existence of the MyD88 signalling pathway (Additional File 1, Figure S2).

**Figure 3:**
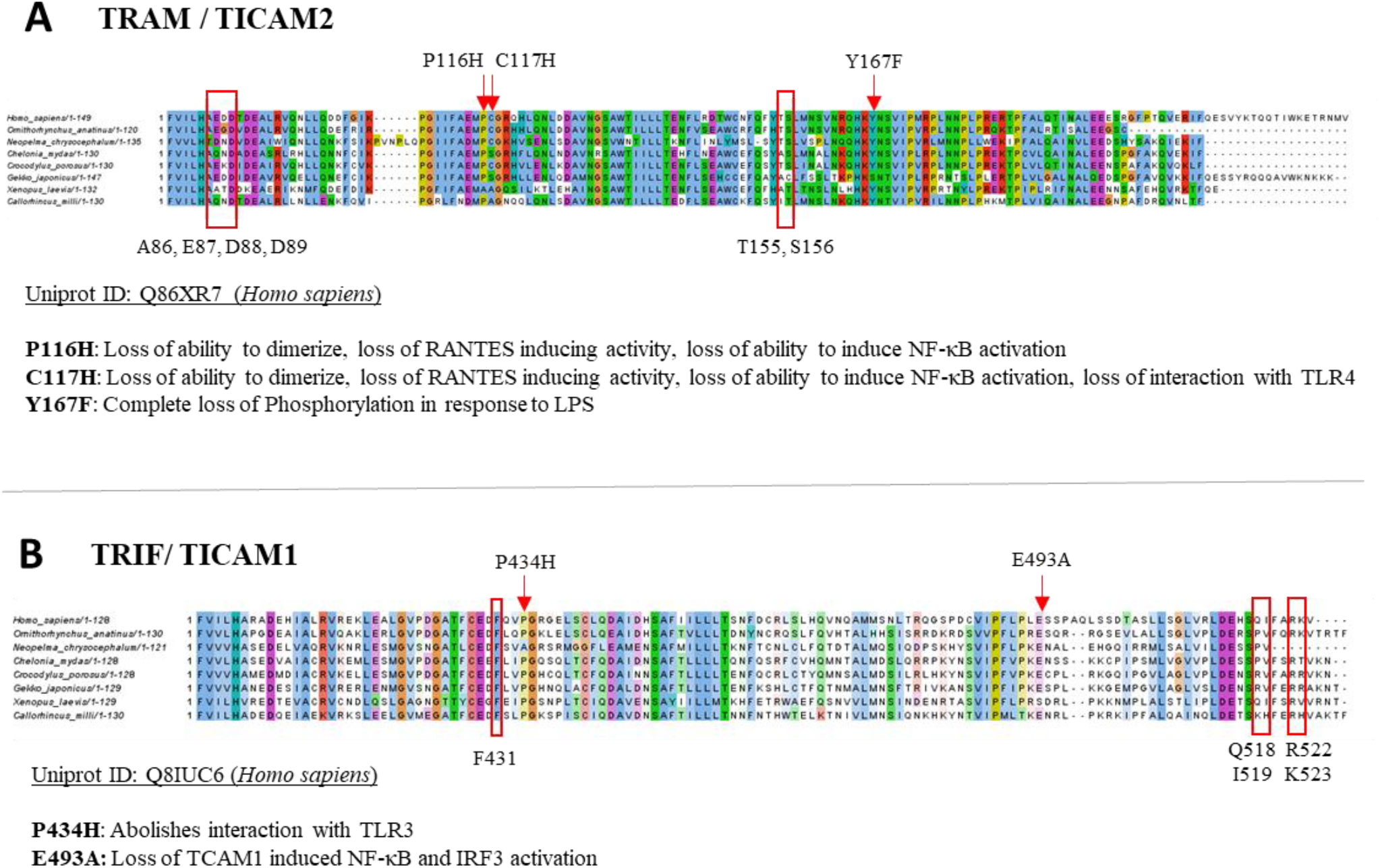
Multiple sequence alignment of **A)** TRAM and **B)** TRIF protein of representative organism. Key functional residues are highlighted in the alignment.

As both the MyD88 dependent and independent pathway are part of TLR4 signalling in humans, we also checked the evidences of these using the Kegg database resources for the representative organism [20].

In TRAM homologues, we found D89 from AEDD site is conserved in all representatives, but the key residues P116 and C117 are both mutated to Alanine in case of *Xenopus laevis*, that may abrogate the TRAM mediated signalling. Also, Y167 is mutated to Serine, in *Gekko japonicus*, although this may still function as phosphorylation site. In case of TRIF orthologs, the F431 is seen well conserved in each representatives, thereby ensuring the interaction of TRIF TIR with TRIF-NTD in autoinhibited stage [12]. Also, the E493 residue is mutated to Serine in case of *Xenopus laevis*. A table listing the key residues from TRAM and TRIF protein is added as Additional File 2, Table S6. Moreover, the conserved motif and the schematic diagram for TLR4 pathway provide deeper insight of these adaptors (Additional File 1, Figure S3 and S4 respectively).

### 3.3 Modelling of human TRAM and TRIF dimers

Interesting conservation and co-evolving residue patterns prompted us to next to model the overall TRAM-TRIF complex. As per the literature we found evidence of TRAM dimer interacting with TRIF monomer [5], where some literature strongly suggests homodimerization of TRIF is important for interferon-β production [21][22].

We had employed several approaches to model TRAM and TRIF dimers in order to model the trimeric and tetrameric models (details in Methodology section). Both dimers were modelled using sequences, docking, homology modelling (HM) and guided docking. The selection of the best TRAM dimer was based on a previous study and that pose where the dimer remained stabilize after molecular dynamics was selected as the best dimer pose.

In case of TRIF dimer similarly we tried blind, guided docking (by BB loop), and homology modelling based on templates of TRAM dimer (as obtained from stabilized MD), TLR6-TIR, TLR10-TIR, and IL1-RAPL TIR structure. We analysed the position of the BB loop and calculated energetics of these models using PPCheck [23]. The models with the energy values are shown in **Figure 4**. The dimer model with Guided docking displays highly stabilizing energy and comparable normalized energy per residues as compared to other models. Moreover, the position of BB loop at the interface and HM_TRAM dimer also seemed more favourable in the guided docking.

**Figure 4:**
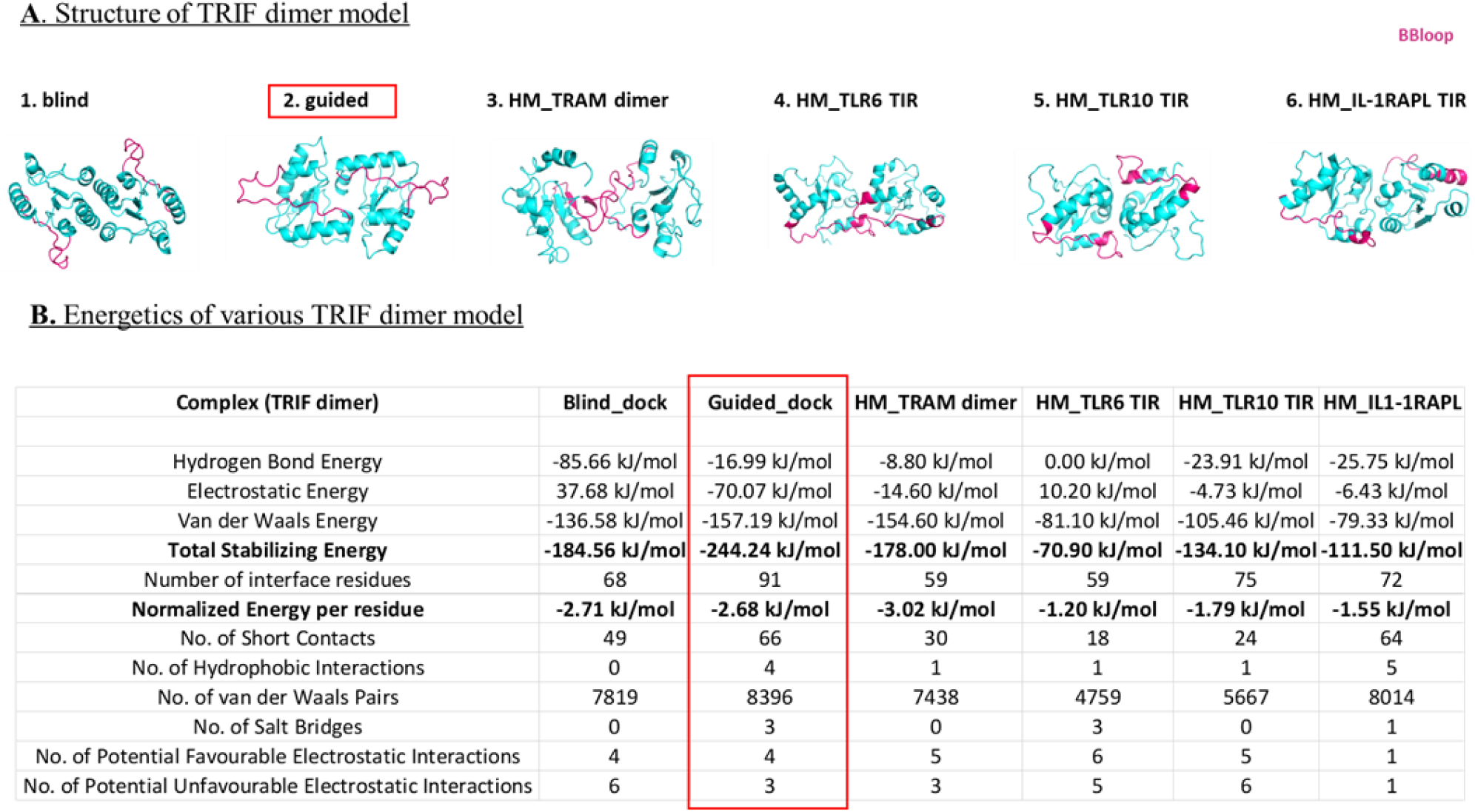
**A)** Dimer model of TRIF using different methods, BBloop represented in pink colour. **B)** Energetics of each dimer model calculated using PPCheck [23].

### 3.4 Modelling of human TRAM TRIF trimer and tetramer complexes

A variety of modelling techniques, including homology modelling, HADDOCK, HADOCK, and Alphafold, were used to construct the trimeric and tetrameric protein complex. All the obtained models were compared based on the energy values calculated using PPcheck, BB loop positions, and key residue positions. We found the model generated using HDOCK docking stands out well in terms of satisfying all the validation parameters. Details of the mode of TRAM TRIF trimeric and tetrameric complexes are provided in Methodology section. We subjected both the modelled complexes to 200ns molecular dynamics simulation and compared the energies of the final frame structure of both the complexes. We found tetrameric complex is more stable than trimeric complex. The details of the energies of each trimeric model and tetrameric models of representative organisms are added in Additional File 2 (Table S2 and S3 respectively). Secondly, we also examined the electrostatic interaction and observed the tetrameric model has complementary potentials at the interface that further explains the higher stability of the complex. The **Figure 5A** and **5B** show the various possible tetrameric models and final frame energies of the tetrameric model. **Figure 5C** also highlights the energy values of the final frame structure of both trimeric and tetrameric model.

**Figure 5:**
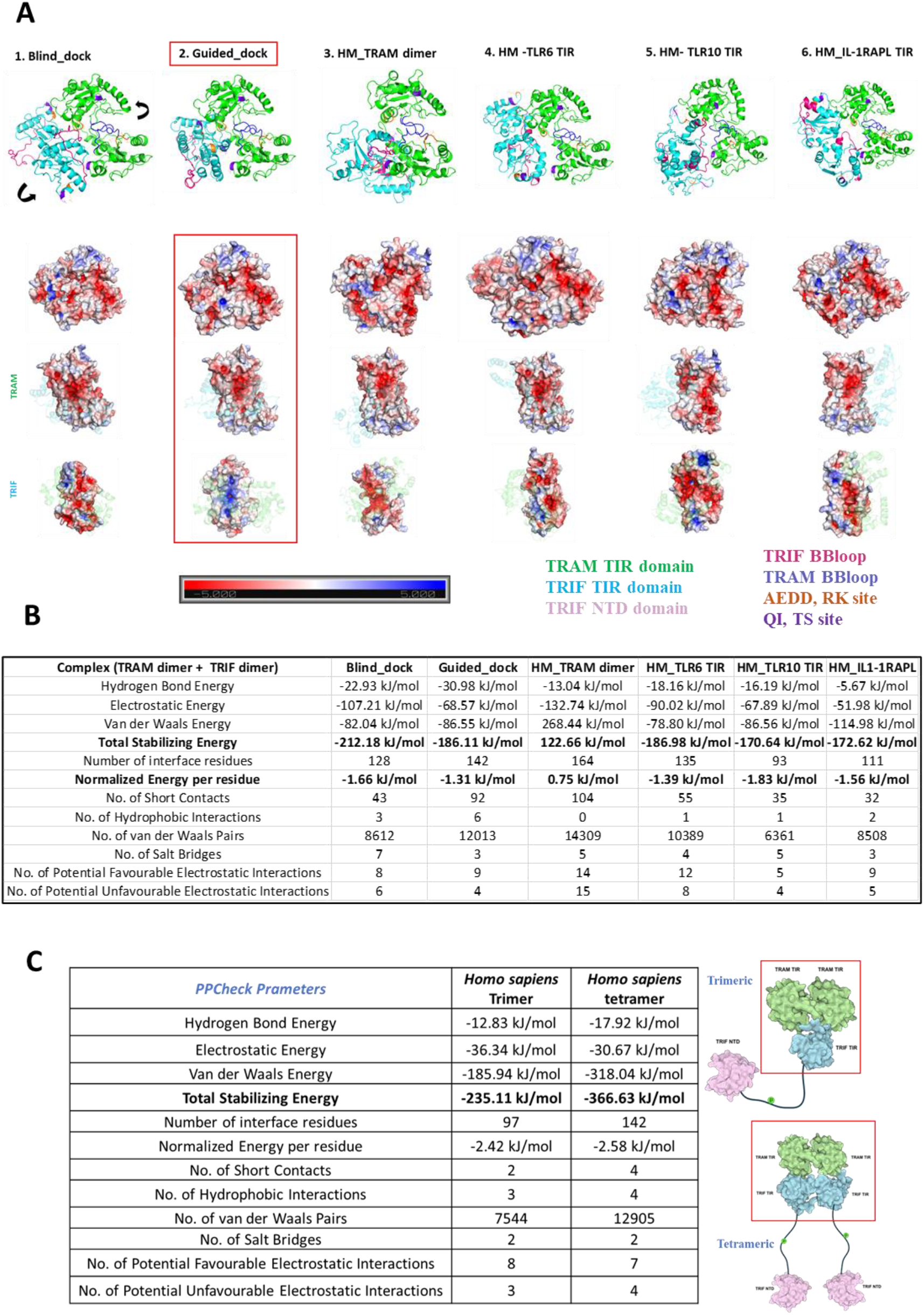
**A)** Multiple approach for tetramer model of *Homo sapiens* TRAM and TRIF dimer. The key residues important for interaction are highted in different colours. The electrostatic potentials of the dimer interface are also shown in the surface diagram. The colour notation of protein in Figure 4A is as mentioned: TRAM TIR dimer in **green**, TRIF TIR dimer in **sky blue**, TRIF BB loop in **pink**, TRAM BB loop in **blue**, AEDD, RK site in **orange** and QI, TS site in **purple. B)** Energetic of various tetrameric model. **C)** Energetic of 200^th^ ns frame molecular dynamic structure trimeric and tetrameric model of TRAM TRIF complex of *Homo sapiens* for the best performed model.

From the **Figure 5A** and **5B** we observe the second model based on the guided docking is performing best in terms of the energy and shows complimentary electrostatic patterns. The **Figure 5C** shows the comparative energy between trimeric and tetrameric model, and we found the total stabilizing energy for the tetrameric model is higher than trimeric model, suggesting it to have higher stability.

### 3.5 Normal mode analysis results (RIN)

Next, we performed normal mode analysis on the complex. We made the complex of orthologs by HDOCK based on key residues based guided docking [24]. The **Figure 6**, below shows the residue interaction plot for representative organism in the trimer and tetramer complex. The residue network plots for trimer and tetramer complex from representative organism is shown in Additional File 1, Figure S5

**Figure 6:**
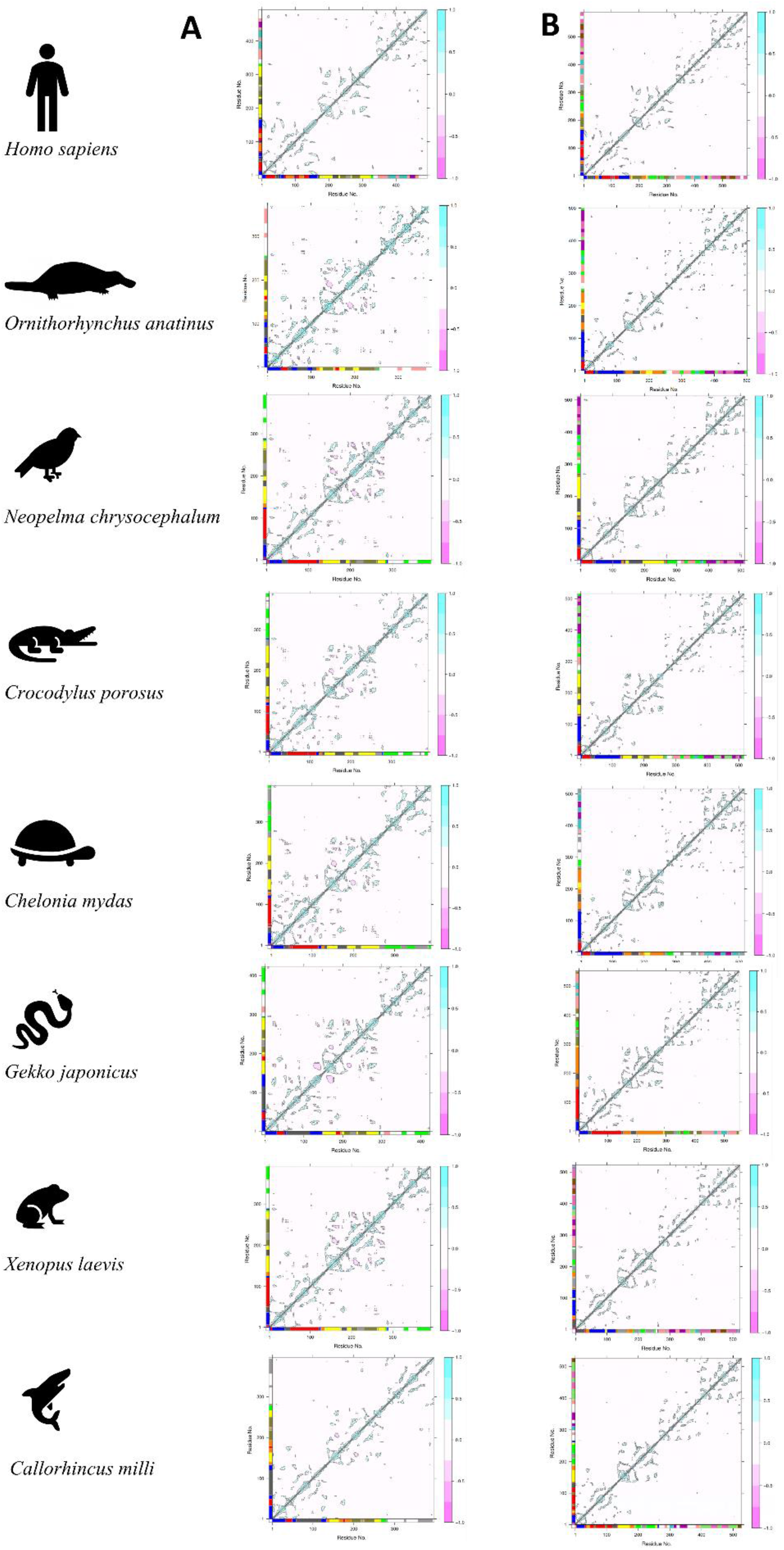
The figure above shows the residue interaction plot of **A**) Trimer complex (TRAM dimer interacting with TRIF monomer) **B)** Tetramer complex (TRAM dimer interacting with TRIF dimer complex).

Here we observe a lot of negative interaction in second chain of TRAM protein of trimer complexes (**Figure 6A**) in all organisms other than *Homo sapiens* (represented by pink colour). That explains the instability of the complex. Whereas in case of tetramer complex (**Figure 6B**) only positive interactions were seen across both the chains of all the organisms (represented by Cyan colour). Meanwhile when we compare the residue clustering pattern in case of trimer (**Additional File 1, Figure S5 A**) we do observe a similar trend across *Neopelma chrysocehalum, Crococodylus porosus, Chelonia mydas, Xenopus laevis* and to some extent in *Gekko japonica*. This is also similar in trend to the known evidences of various adaptor protein involved in TLR4 signalling. Also, from the residue clustering pattern of tetramer (**Additional File 1, Figure S5 B**), we do not observe any cross interaction between TRAM and TRIF protein in case of *Gekko japonica* and *Callorhincus milli* (Australian ghostshark) from the Normal mode analysis. Whereas in other cases there exist connecting clusters between TRAM and TRIF proteins.

### 3.6 Molecular dynamics results (DRN)

The dynamic trend for the modelled protein complexes were ascertained using molecular dynamics (MD) trajectory. A 200ns MD run was compared for the various trimer and tetramer complexes of representative organism. The trajectory was then further used to plot dynamic cross correlation matrix across the complete length of the protein (**Figure 7**).

**Figure 7:**
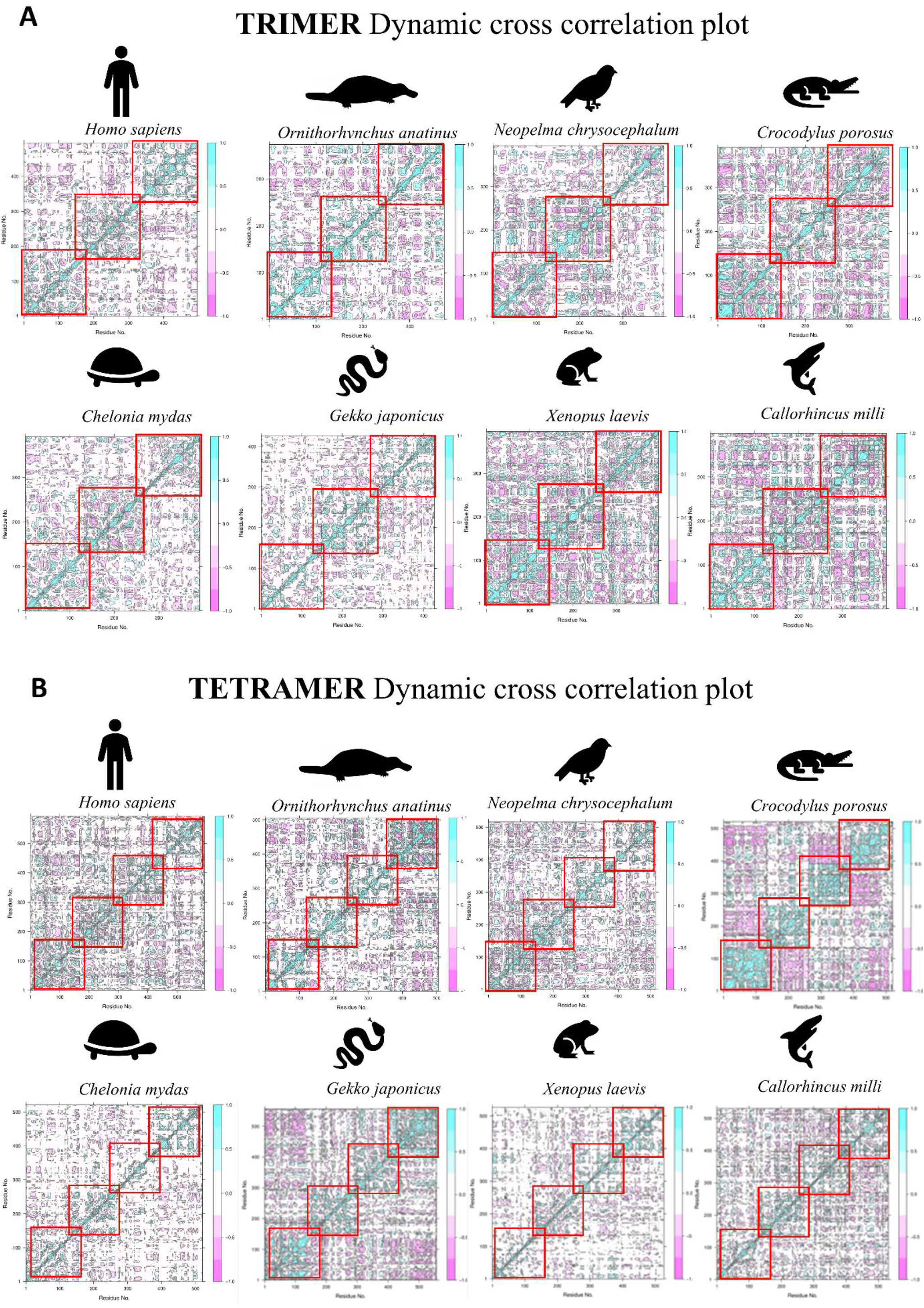
Dynamic cross correlation plot for TRAM TRIF (A) Trimer and (B) Tetramer complexes of representative organisms. The highlighted red boxes represent the intrachain interactions across the MD trajectory. The strength of the positive interaction is shown in cyan colour and negative interaction in pink colour.

While carefully observing the plots we decipher that tetrameric complex shows denser plots thereby stronger interactions. Also, the intensity of positive interaction (cyan colour) is higher in case of tetrameric complex. Thereby it also suggests higher possibility of tetrameric complexes of TRAM and TRIF dimer. Interestingly we observe lesser interaction in case of *Chelonia mydas, Xenopus laevis* that hints towards not much stable complexes in these cases. Additionally, we have also compared the root mean square deviation (RMSD), root mean square fluctuations (RMSF) and conservation pattern of secondary structure of the protein across the trajectory (Additional File 1, Figure S6 and S7 respectively).

Furthermore, using the centrality analysis, and information from the literature we measured several centrality measures of these complexes (Additional File 2, Table S4 and S5). Betweenness centrality being one of the measures of important nodes, was specifically compared for the key hotspot residues (**TRAM**: A86, E87, E88, D89, T155, S156, **TRIF**: Q518, I519, R522, K523), important BB loop residues (TRAM: P116, C117), Phosphorylation site (TRAM:Y167), among different chain of trimer and tetramer complex of representative organism at the respective homologous residue [18][5]. This analysis also highlights the persistence nature of important residues across organisms for trimeric and tetrameric complexes (Additional File 1, Figure S8 and S9 respectively).

## 4. Discussions

### 4.1 Tetrameric complex of TRAM and TRIF TIR domain

Our structural analysis of interactions between TRAM and TRIF from few representative genomes clearly shows that the tetrameric form of the heteromer is more stable than trimeric form, as indicated by the normalised PPCheck energies at the interface. A schematic representing the dimeric tetrameric complex interaction, keeping in mind the key residues involved in interaction is shown below in **Figure 8**.

**Figure 8:**
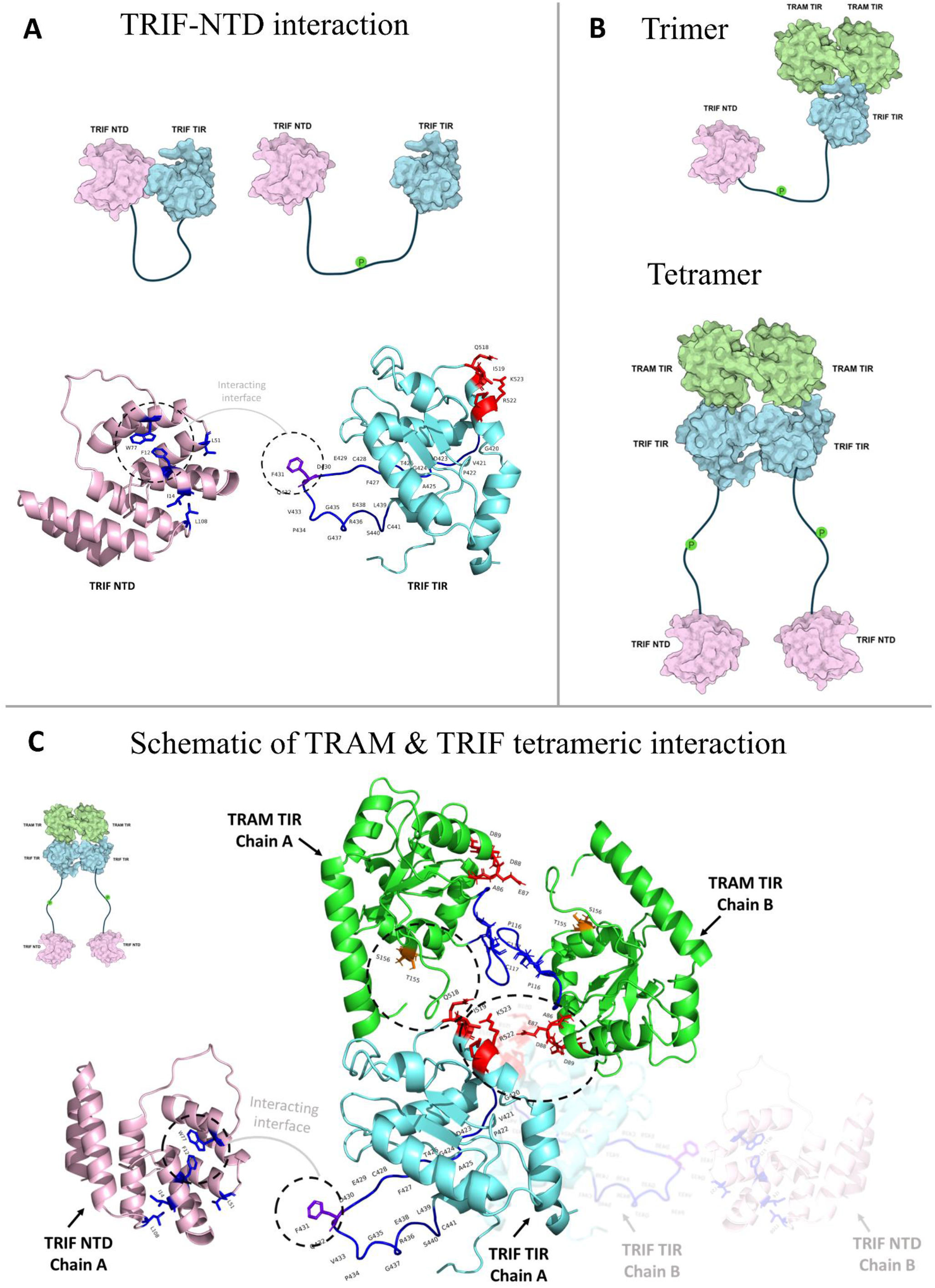
**A)** Schematic of interaction between TRIF N-Terminal domain (NTD) by the F431 of TIR domain [12]. **B)** Schematic showing the trimeric and tetrameric interaction between TRAM and TRIF. It also shows how TRIF-NTD would change orientation post phosphorylation for interaction with IRF3 further downstream. **C)** Schematic showing the TRAM dimer interaction through the BB loop residues, and key residues like AEDD, TS with the QI and RK of TRIF protein in a dimeric way. The TRIF-NTD domain is shown separately, it would be connected to the TRIF TIR by a loop and post phosphorylation separate from TRIF TIR to interact further with IRF3 [25].

### 4.2 Insights from representative organisms

The overall comparison of the various parameters across all the representative organism and surveying the literature associated to the innate immunity we found some interesting facts about the various organisms. In case of *Neopelma chrysocephalum*, it lacks the CD14 protein, that is important for TRAM mediated pathway [26][27]. In case of Avian Toll like receptors, Chickens are widely studied but it lacks the TRAM orthologs like most of the birds [28] [29]. But Aves are known to have developed viral RNA sensing by TRIF mediated TLR3 by producing IFNβ. Furthermore, *Neopelma chrysocephalum* lacks the TRAM Myristoylation motif and TRAF6 binding motif, but has conserved serine in pLxIS motif and have RHIM motif that is important for TRIF induced apoptosis and also contributes towards TRIF induced NF-κB production [30][10]. While observing the *Crocodylus porosus*, it also lacks the CD14 protein, and like Aves viral RNA is similarly sensed by TRIF mediated TLR3 signalling by producing IRF3 and IRF7 [31]. But unlike Aves, *Crocodylus porosus* has both Myristoylation motif and TRAF6 binding motif in TRAM protein and conserved serine in pLxIS motif of TRIF. TRAM in *Chelonia mydas* also lacks the CD14 and has viral RNA sensing by TLR3 mediated signalling [32]. Also, like *Crocodylus porosus*, it has conserved motifs and key residues. In the next taxon representative *Gekko japonicus*, the CD14 protein is also missing. But the presence of alternate pathway recognising LPS was evolving at this level and also positive selection was seen in TLR3 and TLR4 protein of reptiles [33][34]. It also has conserved TRAM and TRIF motifs but lack the interaction as observed through the molecular dynamics study. The amphibian representative *Xenopus laevis* similarly lacks the CD14 and also the Myristoylation motif on TRAM, that is crucial for TRAM localization to plasma membrane [9]. Besides this while studying the ancestor most organism *Callorhincus milli*, TRF6 binding motif of TRAM protein was missing, but Myristoylation motif of TRAM and TRIF’s motif remain conserved. Although the interaction study shows distinct cluster for TRAM and TRIF of this taxa, and the absence of key residue suggests the TRAM TRIF mediating signalling might be evolving [35]. A schematic representing the conserved motif is added in Additional File 1, Figure S3.

Even before the Chondrichthyes (*Callorhincus milli)*, the ancestor of TRAM and TRIF proteins were observed in Leptocardii (*Branchiostoma belcheri*) also known as amphioxus [36]. The evidence of the emergence of Myd88-independent pathway in amphioxus by discovery of a novel TIR adaptor referred as bbtTICAM had widen the scope of TRAM TRIF mediated pathway in invertebrates. Even though the bbtTICAM activates NF-kB in MyD88 independent manner by the interaction of TRIF’s RHIM domain, but it fails to induce the production of TRAM TRIF mediated type I interferons [37]. As per the computational analysis of the common ancestor of both the adaptor protein TRAM and TRIF we observe *Callorhincus milli* lacks the key residues of TRAM that may similarly hinder in the type I interferons signalling. The conservation of key functional residues in TRIF would suggest the presence of MyD88-independent NF-kB activation by the TRIF’s RHIM domain involving TLR3.

## 5. Conclusions

Sequence and structural patterns of TIR domains of TLR4 adaptors (TRAM and TRIF protein) among representative orthologs across tree of life [4] provides bird’s eye of view of the evolutionary trajectory of Toll-like receptors. Our modelling approach has been an attempt to decipher the possible interaction mechanism, and while comparing the dynamics we also observe the stability of tetrameric complexes being higher than the trimeric ones. The study has also focussed on key residues to examine if their interactions are persistent across the simulations. To understand the however, it is imperative to study and focus on such pathways to get deeper insights and gain insights on complete functional TRAM mediated signalling pathways across various taxa and evolution of innate immunity in non-mammals.

In this paper, we have employed modelling and molecular dynamics of TIR assemblies in the TLR4 pathway to show that the conservation of functional motifs or the cross-talk between them might be affected in the primitive TIR adaptor domains. Hence, either the Myd88-dependent pathway or the TLR3-TRIF mediated pathways maybe operating in the ancient TRAM adaptors, like Amphioxus and ghostshark.

Besides TLRs, the presence of other pattern recognition receptors (PRRs) in humans are also accountable for the innate immunity. These include the nucleotide-binding domain leucine-rich repeat/NOD-like receptors (NLRs), RIG-I-like receptors (RLRs), and C-type lectin receptors (CLRs). Their role in the identification of various ligands from different viruses, bacteria, fungus, and other pathogens makes them an essential first line of defence [38]. Evolutionary study of these PRRs across various taxa will shed more light on the defence mechanism of immune pathway of various organisms.

## 6. Materials and methods

### 6.1 Homologous protein analysis

The ortholog hits were selected based on specific and conserved positions of amino acid for the TIR_2 domain of TRAM (PDB ID: 2M1W) and TRIF (PDB ID: 2M1X) [4]. These ortholog sequences were searched using 30 mammalian query sequences across a non-redundant database by genome-wide search method using CS-BLAST methods. These hits were filtered based on query coverage cutoff (>50%), and percentage identity cutoff (>30 %). The sequences were also categorized based on conserved motifs and domain patterns. The detailed study on these sequences is presented in one of the previous studies from our lab [4]. On these sequence sets, **CONSURF** was then used to visualize conservation in protein sequence amongst evolutionary orthologs. These mappings were done wrt TIR_2 domain on PDB structure (ID: 2M1W and ID:2M1X), and in case of mutations (2M1W: H117C, 2M1X: P434H) they were reverted to wild type [15]. These sequences were further analysed using the **Evolutionary Trace** (ET) method to compute the relative rank of functional and structural position among protein homologs [14]. Further, **visualCMAT** was used to analyse co-evolving residues [16].

### 6.2 Protein Stability Analysis

The coevolving residue pairs that belong to highly conserved category from CONSURF analysis (score > 6) were further analysed. These pairs were first checked for the frequency (>1%) of their occurrences in the orthologs. The corresponding amino acid changes were then incorporated using FoldX 5.0 [19]. **RepairPDB** command was used initially on TRIF and TRAM TIR domain structure (2M1W; H117C, 2M1X; P434H) to identify residues with bad torsion angles, Van der Waals clashes or total energy and repairs them. The other parameters include pH=7, temperature=298K, ionStrength=0.05M and vdwDesign=2. After repairing the structures, **BuildModel** command was used to mutate the residues from coevolving pairs to differently observed combinations. These runs were iterated for five times with same parameters as in RepairPDB and then the Average score for **ΔΔG kcal/mol** (ΔGmut-ΔGwt) was calculated. The results were binned in different categories based on the ΔΔG values as follows: *highly stabilising* (ΔΔG < −1.84 kcal/mol); *stabilising* (−1.84 kcal/mol ≤ ΔΔG < −0.92 kcal/mol); *slightly stabilising* (−0.92 kcal/mol ≤ ΔΔG < −0.46 kcal/mol); *neutral* (−0.46 kcal/mol < ΔΔG ≤ +0.46 kcal/mol); *slightly destabilising* (+0.46 kcal/mol < ΔΔG ≤ +0.92 kcal/mol); *destabilising* (+0.92 kcal/mol < ΔΔG ≤ +1.84 kcal/mol); *highly destabilising* (ΔΔG > +1.84 kcal/mol).

### 6.3 Protein Modelling and docking of trimer complex

We used multiple approaches for modelling proteins dimer, trimer, and tetramer complex. In a previous study from our lab, we had established the dimer model of TRAM protein using various parameters [39]. In one of the approaches, we used the TRAM MD stabilized structure as our starting dimer model and use protein docking algorithm to model the TRAM dimer and TRIF complex. We also used HADDOCK and HDOCK webserver to dock the models by both blind and guided docking [40] [41] [24]. Following the current trend in the field we also used AlphaFold colab notebook that used a slightly modified version of AlphaFold v2.3.2. This is a template independent modelling, that is trained on the BFD database[42]. Inclusive of all we used six different approaches, and modelled TRAM-TRIF TRIMER complex as reported in previous literature [5]. Here in approach 1, we used AlphaFold modelled TIR of TRAM and TRIF and docked using the known interacting residues from literature (AEDD & TS site of TRAM and QI & RK site of TRIF) using HADDOCK. In approach 2, we used MD stabilised final frame structure (200^th^ ns frame) of TRAM dimer and performed bling docking using HDOCK. In approach 3, we used structure modelling of TRAM and TRIF from their respective sequence followed by blind docking using HDOCK. In approach 4 we completely used the AlphaFold to build the multimer complex. In approach 5 and 6 somewhat similar to the approach 4 we used MD stabilised final frame structure of TRAM dimer and did guided docking (AEDD & TS site of TRAM and QI & RK site of TRIF [5]) using HADDOCK and HDOCK method respectively.

Latter we analysed these complexes based on the positioning of the key residues, positioning of the BB loop, and energy calculations at the interface (calculated using PPCheck [23]). We found substantiated evidence from the literature that the HDOCK is more efficient as compared to other methods and also based on validation parameters it stands as the best methods [43]. Thereby we used the 6^th^ approach model of TRAM TRIF timer complex for further analysis. The structure of the trimeric complex and its energies is shown in Additional File 1, Figure S10.

### 6.4 Protein Modelling and docking of tetramer complex

There is limited literature evidence of TRIF dimer interacting with TRAM dimer [25] [21]. To investigate this possibility further, we modelled TRIF dimer using multiple approaches. We used HDOCK for blind, and guided docking of TRIF dimer (dimer along BB loop, as observed in most TIR domain interactions). In other approach we used our most stabilised structure of TRAM dimer as a template and performed homology modelling for TRIF dimer. We also used template-based TRIF dimer modelling using the available structure in PDB (TLR6 (PDB ID: 4OM7) TLR10 (PDB ID: 2J67) and IL-1RAPL (PDB ID: 1T3G).

Further, we used these six TRIF dimer models and established the TRAM and TRIF tetramer complex. All these combinations were further compared based on the energetics values and significant residue positions. We also examinedthe electrostatics at the dimeric interface using APBS plugin of PyMol software [44].

### 6.5 Selection of representative organism

Homology modelling was performed for some of the representative organisms from different taxa across the tree of life using Modeller [45]. In a previous study, we have traced the evolution of TRAM and TRIF protein from the oldest ancestors [4]. We analysed the domain architecture and gained deep insights on the residue conservation pattern. From the corresponding study, we selected representative organisms from different taxa, such that it consists of well annotated TRAM and TRIF TIR domains as it is crucial for signalling. *Callorhincus milli* (Chondrichthyes) was the oldest ancestor with both TRAM and TRIF TIR domains. *Xenopus laevis* (Amphibians), *Chelonia mydas* (Cryptodira), *Crococodylus porosus* (Crocodylia), *Gekko japonica* (Bifurcata), *Neopelma chrysocehalum* (Aves), and *Ornithorhynchus anatinus* (Mammalia) was chosen as representatives of each taxon.

### 6.6 Normal mode analysis

We used the Bio3d package, in R for comparative analysis of protein structures [46]. We performed the Normal mode analysis for capturing the large-scale molecular motions of the proteins. To obtain a detailed understanding at all atom level, we performed all-atom normal mode analysis (ENM) [47]. Hessian matrix was calculated for the protein complex and lowest frequencies modes were observed [48]. The initial six modes represent the trivial modes with zero frequency corresponding to the rigid-body rotation and translation. We performed dynamic cross correlation analysis, plotted residue interaction network and finally network analysis (using cna function). We observed several numbers of network communities in each complex. Later, fluctuation and deformation analyses were performed as well. These measures provide us amplitude of absolute atomic motions and the amount of local flexibility in the protein structure, respectively.

### 6.7 Molecular dynamics simulations

The complexes were also subjected to molecular dynamics (MD) simulation to obtain a detailed understanding of the all-atom movements. Initially, we prepared the protein structure, using protein preparation wizard of Maestro from Schrodinger suite [49]. The pre-processing was performed to cap the termini and by filling the missing residues, if any. Then H-bond assignments were optimized, water molecules were deleted from the complex and structure was overall minimized. The system builder option was then used to prepare the system. TIP4P explicit solvent model was chosen, and a orthorhombic box was defined with minimum volume size [50]. The OPLS4 force field was selected and the whole system was neutralized by adding equivalent number of ions [51]. Additionally, to mimic the physiological conditions, the salt concentration was maintained with 150mM NaCl. The complex was further relaxed and a simulation of 200ns was run at NPT conditions using Desmond from Schrodinger suite [52]. The trajectory was further converted and used for dynamic cross correlation analysis using the Bio3d package [46].

### 6.8 Residue Network Analysis

The calculated MD trajectory was further used to perform dynamic residue network (DRN) analysis using multiple trajectory frames. For the above-mentioned analysis, residue network graph was constructed using C-alpha atoms as nodes, that were connected to each other by edges with a defined cut-off distance of 6.7 Å for each protein residue pair. Several DRN metrics like Betweenness Centrality (BC), Average Shortest Path Length (L), Closeness Centrality (CC), Eccentricity (ECC), Degree Centrality (DC), Eigencentrality (EC), Katz centrality (KC) and PageRank (PR) were calculated using the MDM-TASK-web platform [53].

## 8. Ethics approval and consent to participate

Not applicable

## 10. Author Contributions

Shailya Verma: Formal analysis; visualization; data curation; writing – original draft. R. Sowdhamini: Conceptualization; formal analysis; writing – review and editing; supervision; funding acquisition; resources; project administration

## 11. Authors’ information (optional)

## 12. Acknowledgments

The authors would like to thank NCBS (TIFR) for infrastructural facilities. They would like to thank Dr Vinoth Kumar, Dr Praveen Vemula and CAPS lab members for useful discussions. RS acknowledges funding and support provided by JC Bose Fellowship (JBR/2021/000006) from Science and Engineering Research Board, India and Bioinformatics Centre Grant funded by Department of Biotechnology, India (BT/PR40187/BTIS/137/9/2021). RS would also like to thank Institute of Bioinformatics and Applied Biotechnology for the funding through her Mazumdar-Shaw Chair in Computational Biology (IBAB/MSCB/182/2022).

## 13. Data Availability statement

The authors declare that [the/all other] data supporting the findings of this study are available within the article [and its supplementary information files].

## 14. Additional File

### Additional File 1

1. **Figure S1:** The details of the free energy change due to coevolving mutation in TRAM and TRIF
2. **Figure S2:** The conservation pattern in MyD88 and TIRAP protein to validate the existence of the MyD88 signalling pathway
3. **Figure S3:** The phylogeny showing the conserved motifs and domain architectures of representative sequences from TRAM and TRIF protein
4. **Figure S4:** The schematic diagram for the TLR4 pathway of representative organisms.
5. **Figure S5:** The residue network plots for trimer and tetramer complex from the representative organisms.
6. **Figure S6:** The root mean square deviation (RMSD) of the trimeric and tetrameric protein across the trajectory
7. **Figure S7:** The root mean square fluctuations (RMSF) of the trimeric and tetrameric protein across the trajectory
8. **Figure S8:** The centrality measures highlighting the persistence nature of important residues across organisms for trimeric complexes
9. **Figure S9:** The centrality measures highlighting the persistence nature of important residues across organisms for tetrameric complexes
10. **Figure S10:** The structure of the various trimeric complex models and its energies

### Additional File 2

1. **Table S1:** Frequency of co-evolving pairs A) TRIF-TIR, B) TRAM-TIR
2. **Table S2:** Energies of Trimer model of representative organisms
3. **Table S3:** Energies of Tetramer model of representative organisms
4. **Table S4:** Centrality measures of Trimeric model of representative organisms
5. **Table S5:** Centrality measures of Tetrameric model of representative organisms
6. **Table S6:** key residues of TRAM and TRIF protein

