## Additional File 1 for "Sequence and structural analysis of adaptors of Toll-like receptor 4 sheds light on the evolutionary trajectory and functional emergence"

**A**

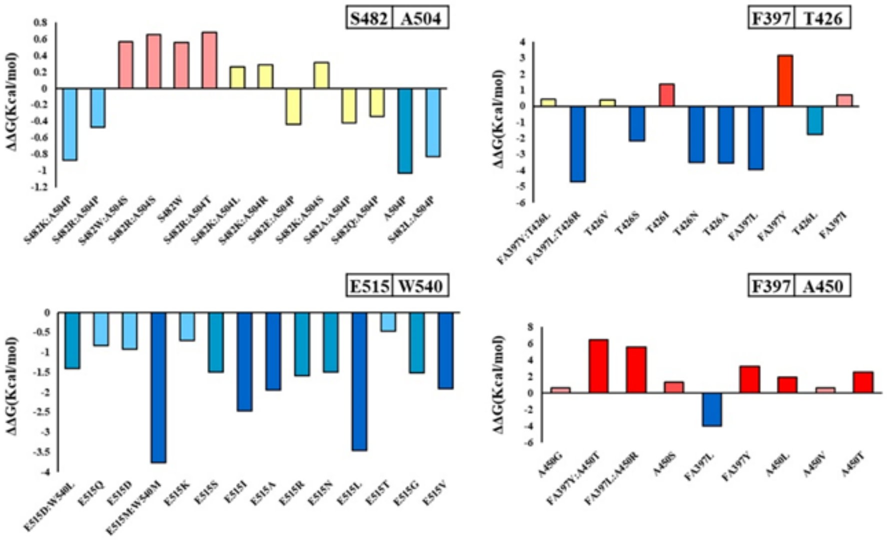

| Co-evolving pairs | | MIp | MIc | Zp | Zc |
|---|---|---|---|---|---|
| S482 | A504 | 0.196 | 0.182 | 4.438 | 4.678 |
| E515 | W540 | 0.172 | 0.169 | 3.906 | 4.381 |
| F397 | T426 | 0.159 | 0.155 | 3.608 | 4.037 |
| F397 | A450 | 0.157 | 0.153 | 3.565 | 3.986 |

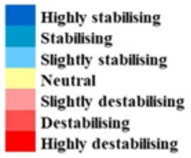

**B**

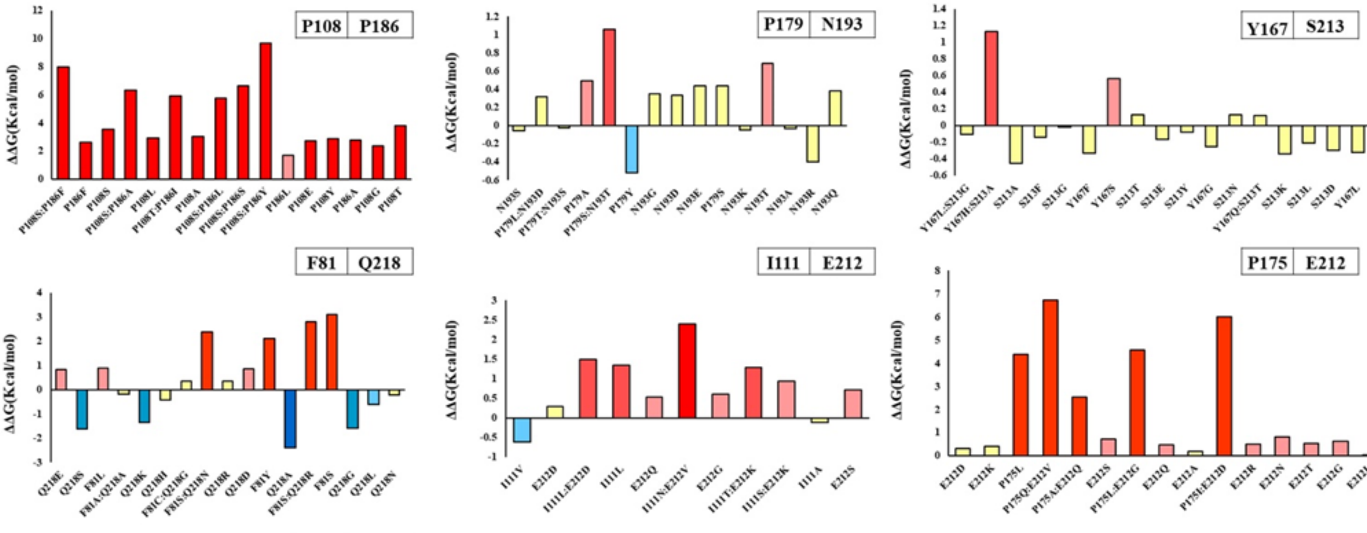

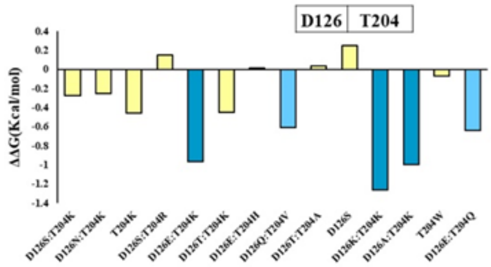

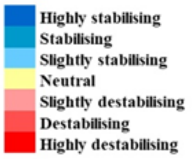

| Co-evolving pairs | | MIp | MIc | Zp | Zc |
|---|---|---|---|---|---|
| P108 | P186 | 0.127 | 0.117 | 4.031 | 4.162 |
| P179 | N193 | 0.115 | 0.109 | 3.669 | 3.898 |
| Y167 | S213 | 0.113 | 0.108 | 3.598 | 3.873 |
| F81 | Q218 | 0.111 | 0.106 | 3.546 | 3.773 |
| I111 | E212 | 0.111 | 0.105 | 3.536 | 3.771 |
| P175 | E212 | 0.111 | 0.102 | 3.528 | 3.664 |
| D126 | T204 | 0.127 | 0.1 | 4.036 | 3.564 |

**Figure S1**: **A**) shows the ΔΔG values from different coevolving pairs of TRIF protein and **B**) for TRAM protein. The free energy change value is shown by different colours based on stabilising or destabilising nature. Table shows the result from VisualCMAT of coevolving pairs, where MIp and MIc are depicted from mutual information-based statistics and Zp, Zc are the Z score that should be > 3.5 for further consideration. The colour of the bar plot shows the nature of the mutation and is binned into category based on different scores (kcal/mol) as follows: highly stabilising ($\Delta\Delta G < -1.84$); stabilising ($-1.84 \le \Delta\Delta G < -0.92$); slightly stabilising ($-0.92 \le \Delta\Delta G < -0.46$); neutral ($-0.46 < \Delta\Delta G \le +0.46$); slightly destabilising ($+0.46 < \Delta\Delta G \le +0.92$); destabilising ($+0.92 < \Delta\Delta G \le +1.84$); highly destabilising ($\Delta\Delta G > +1.84$).

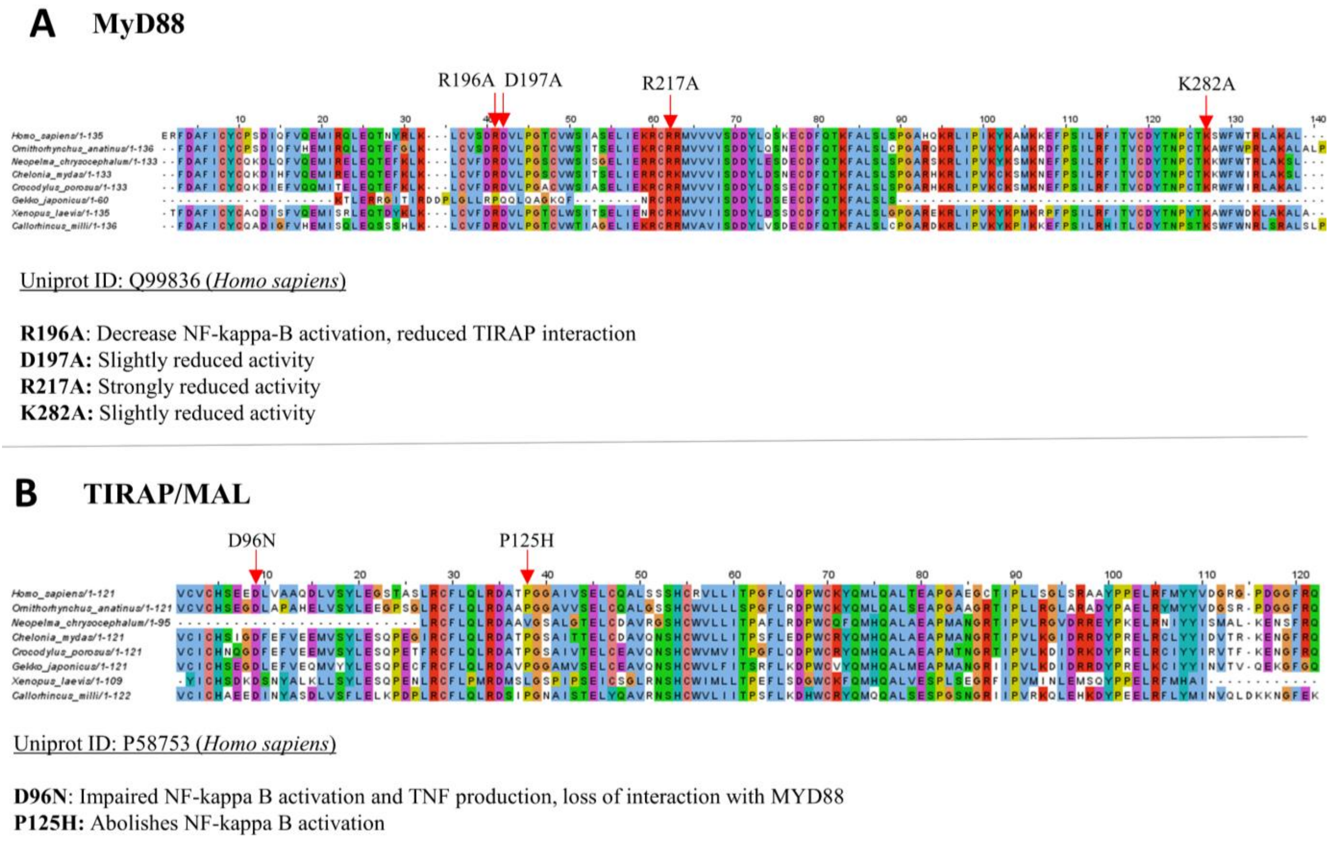

**Figure S2: A)** Sequence alignment of MyD88 protein from representative organisms. The key residues as per *Homo sapiens* Uniprot entry (ID: Q99836) are highlighted with mutational effects. **B)** Sequence alignment of TIRAP protein from representative organisms. The key residues as per *Homo sapiens* Uniprot entry (ID: P58753) are highlighted with mutational effects.

**Representative sequences of TRAM protein**

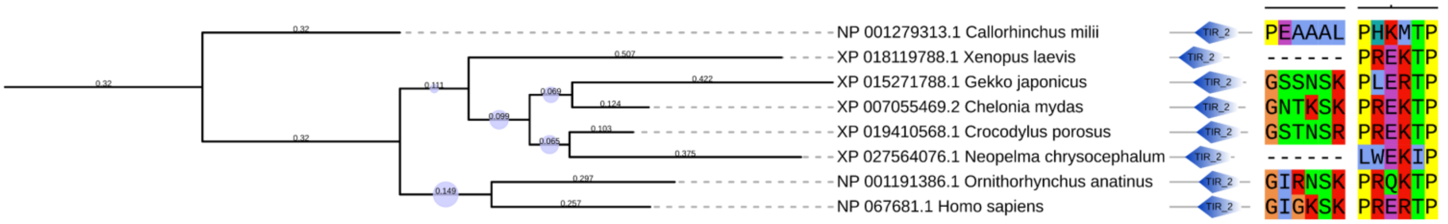

- N-Myristoylation site has a consensus motif sequence of [G{EDRKHPFYW}xx[STAGCN]{P}]
- TRAF6 binding motif [PxExxP]

**Representative sequences of TRIF protein**

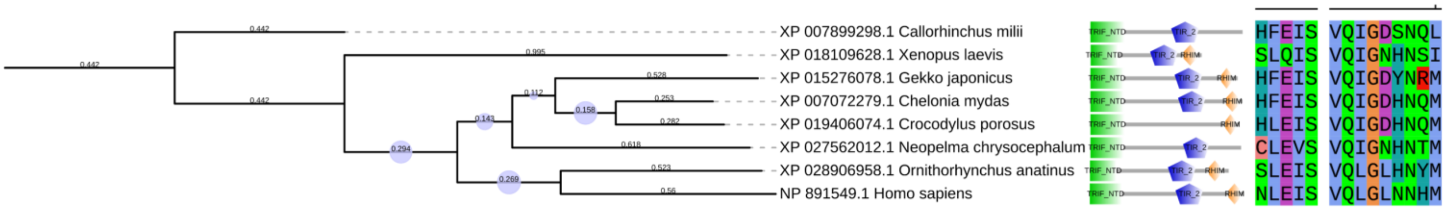

- pLxIS motif ([[NDQEKR]LxIS])
- RHIM interacting motif ( [[IV]Q[ILV]GxxNx[MLI]])

**Figure S3:** The phylogeny showing the conserved motifs and domain architectures of representative sequences from **A) TRAM** and **B) TRIF** protein

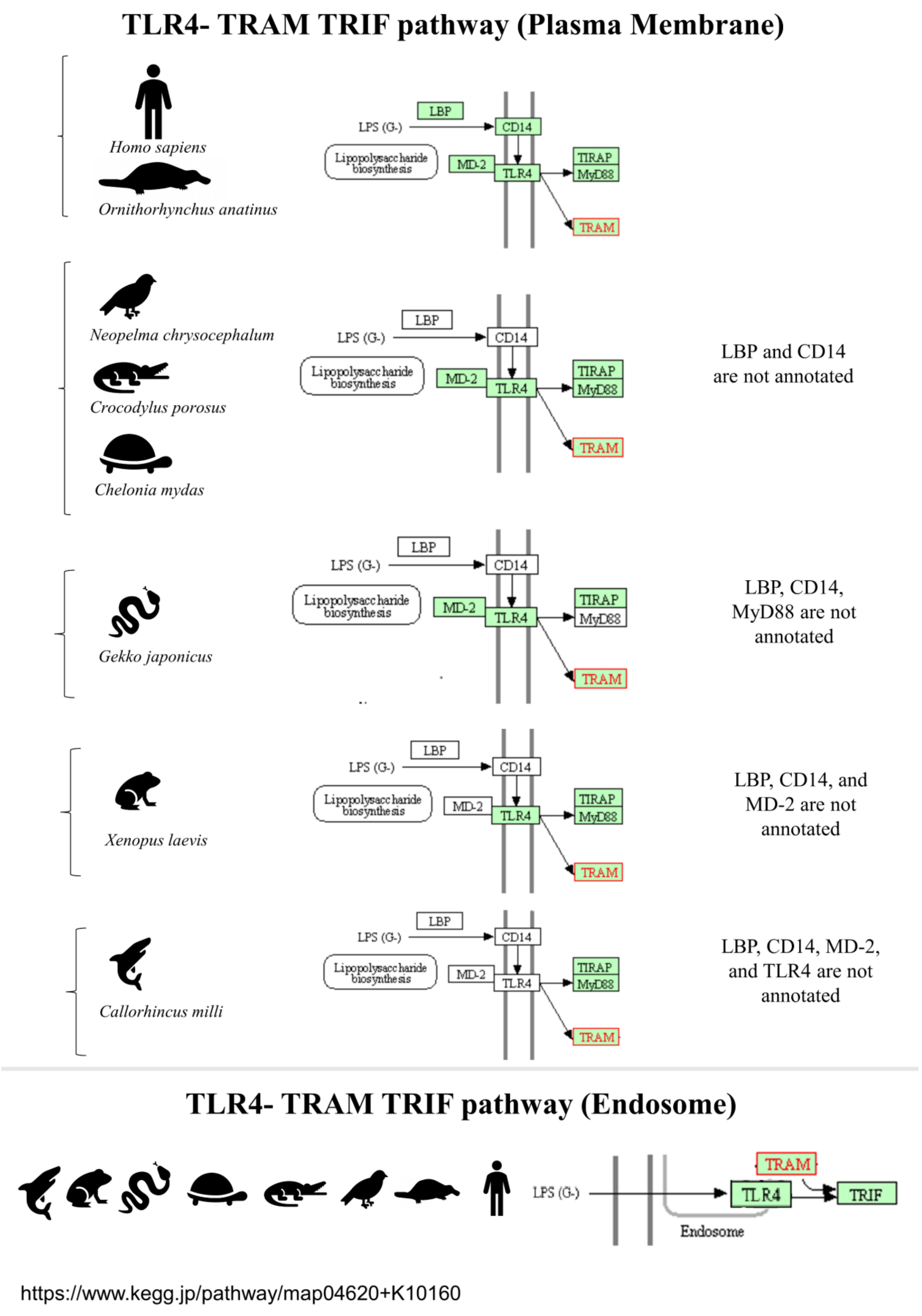

**Figure S4:** The schematic diagram for the TLR4 pathway of representative organisms. The pathways are taken from the Kegg database.

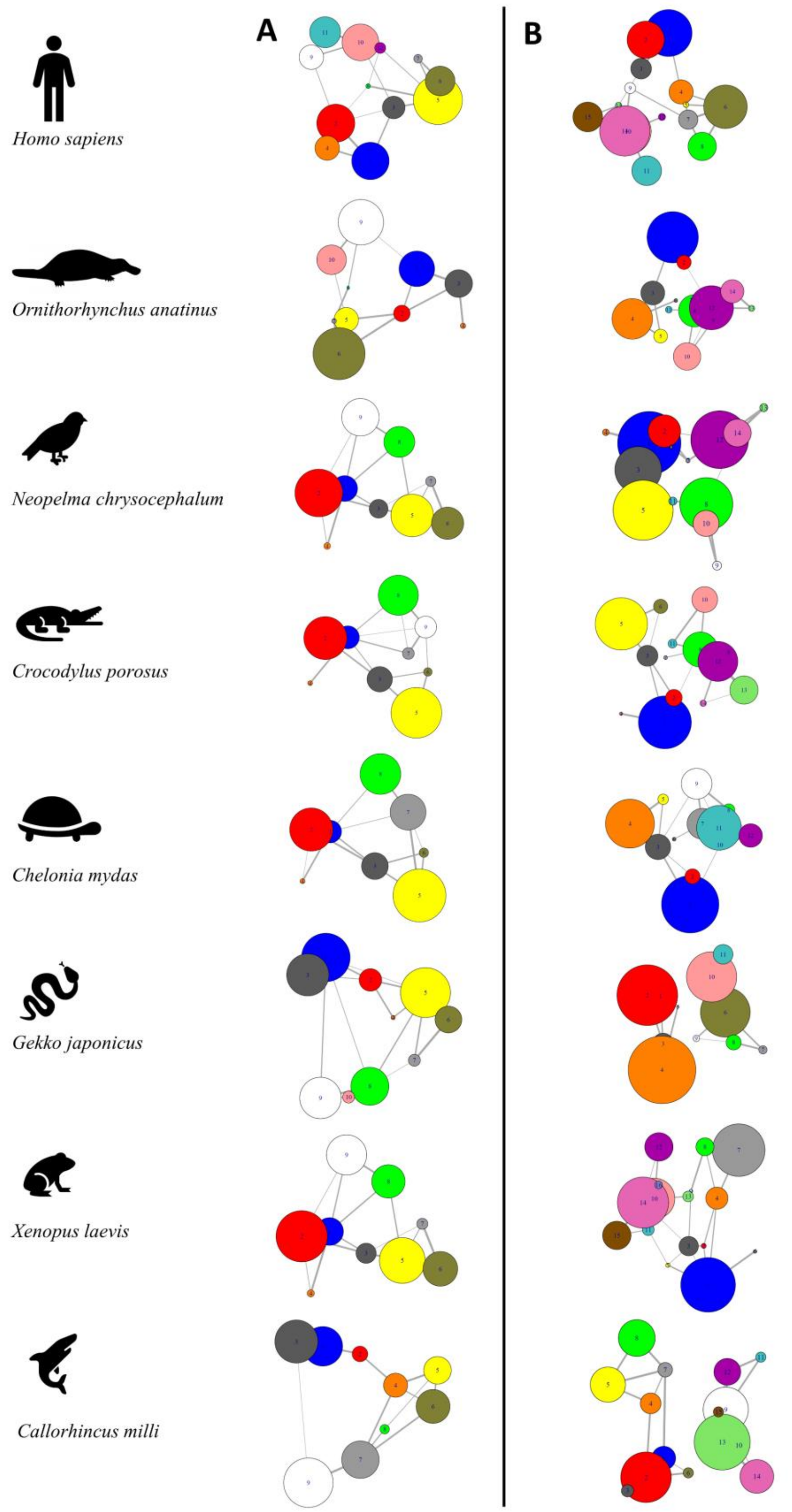

**Figure S5:** The residue network plots for trimer and tetramer complex from the representative organisms.

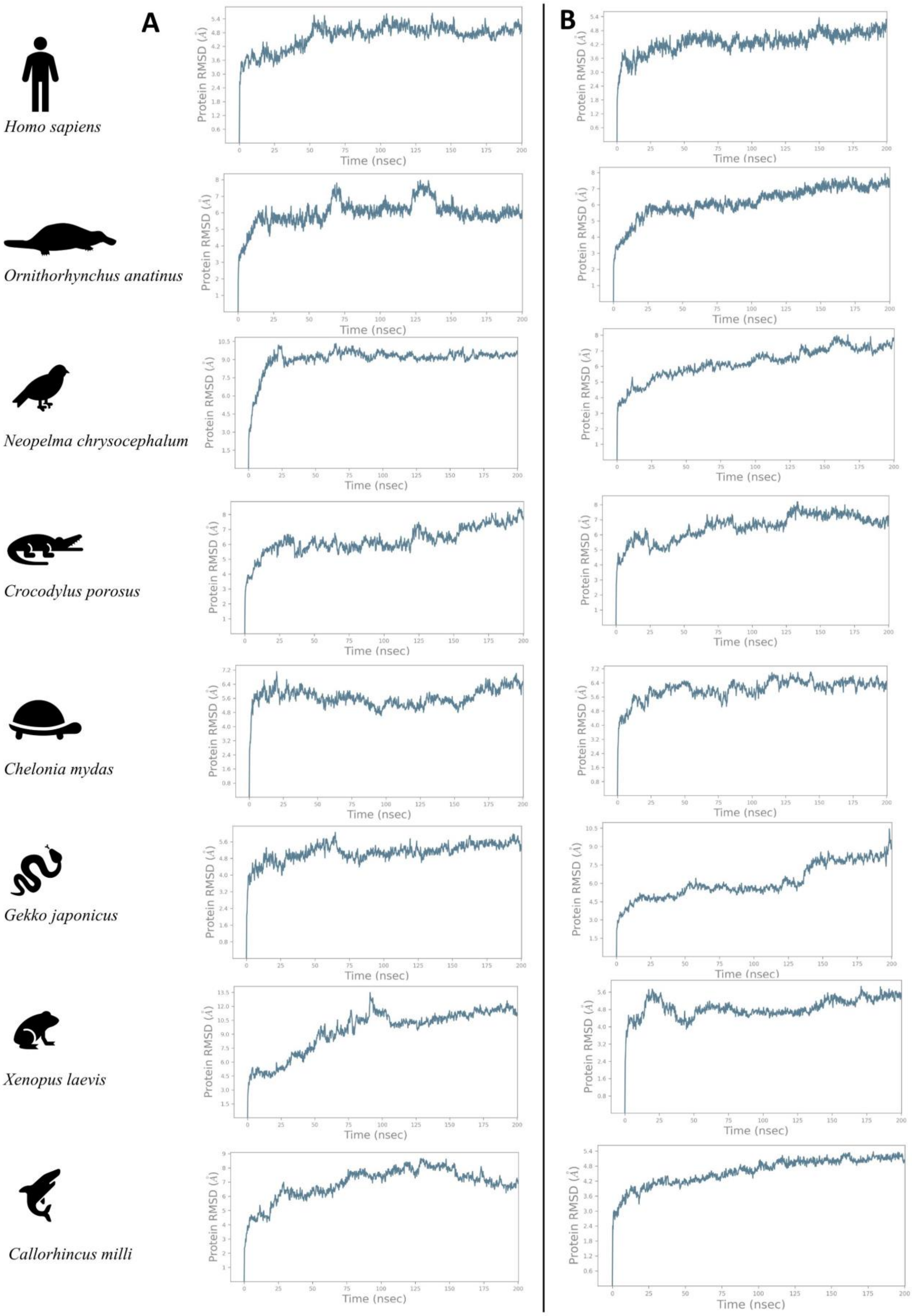

**Figure S6:** The root mean square deviation (RMSD) of the trimeric and tetrameric protein across the trajectory

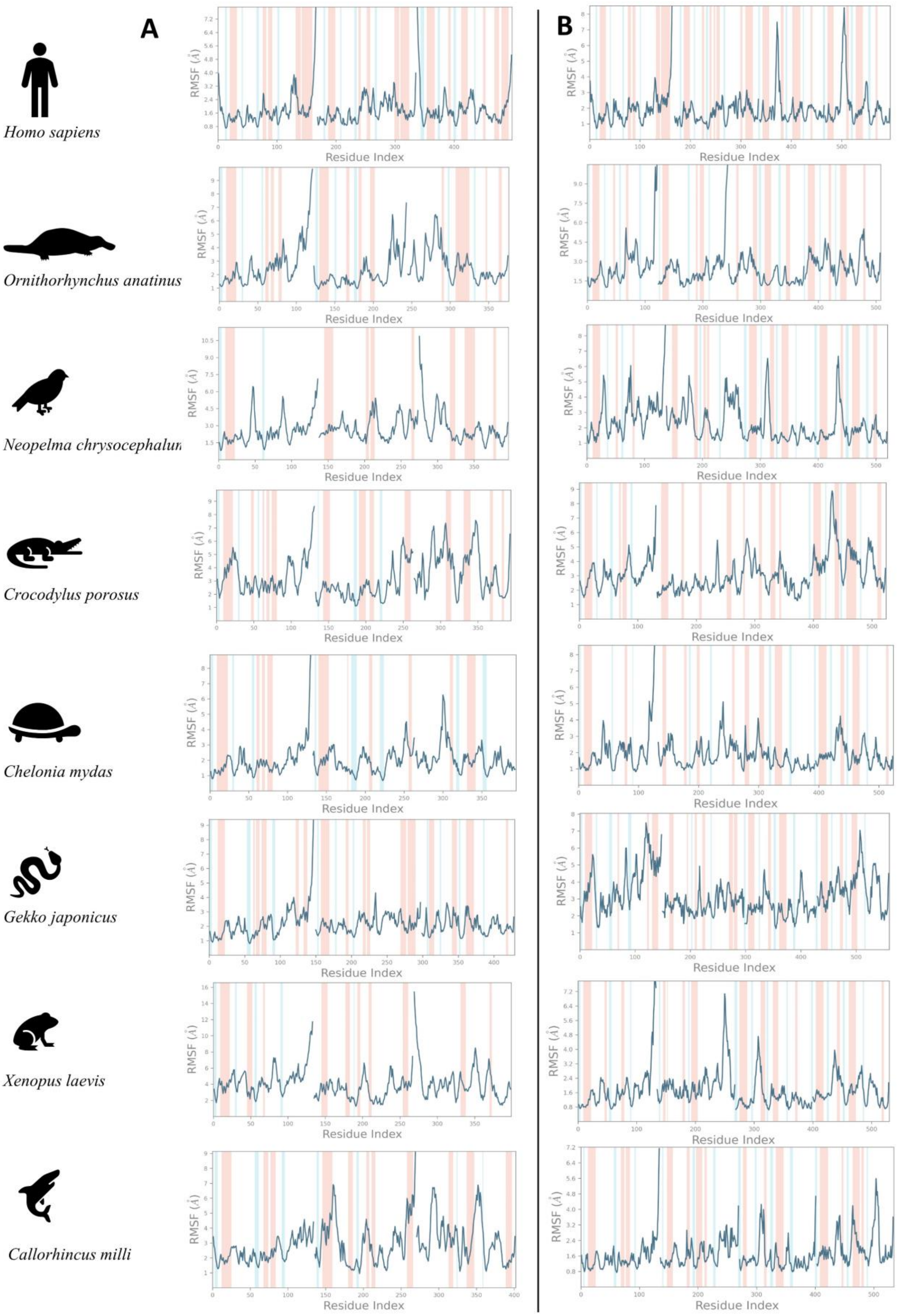

**Figure S7:** The root mean square fluctuations (RMSF) of the trimeric and tetrameric protein across the trajectory

| residue no. | residue | *H. sapiens* | *O. anatinus* | *N. chrysocephalum* | *C. porosus* | *C. mydas* | *G. japonicus* | *X. laevis* | *C. milli* |
|---|---|---|---|---|---|---|---|---|---|
| **TRAM-TIR** | **Chain A** | | | | | | | | |
| 86 | **A** | 0.439688751 | 0.44773378 | 0.137988952 | 0.639636623 | 0.44529373 | 0.345054242 | 0.3810811 | 0.536532 |
| 87 | **E** | 0.253090049 | 0.19293598 | 0.084408276 | 0.574034022 | 0.16040455 | 0.276362794 | 0.396633 | 0.453406 |
| 88 | **D** | 1 | 0.750284051 | 0.039434721 | 0.922784501 | 1 | 0.070487845 | 0.7255782 | 0.869531 |
| 89 | **D** | 0.66600622 | 0.369039014 | 0.327888492 | 0.433671258 | 0.62525695 | 0.572358201 | 0.7335865 | 0.458184 |
| 116 | **P** | 0.585223444 | 0.44698685 | 0.809881421 | 0.453850972 | 0.65285883 | 0.453486624 | 0.0383417 | 0.064726 |
| 117 | **C** | 0.401509113 | 0.529615054 | 0.272381463 | 0.44356788 | 0.36080966 | 0.292136081 | 0.1468775 | 0.129874 |
| 155 | **T** | 0.010452791 | 0.004604687 | 0.021131689 | 0.019597128 | 0.01067756 | 0.002350918 | 0.0263856 | 0.006992 |
| 156 | **S** | 0.09045416 | 0.074727846 | 0.106809406 | 0.138095733 | 0.08305901 | 0.043020659 | 0.2441903 | 0.046079 |
| 167 | **Y** | 0.054964797 | 0.029994565 | 0.097748975 | 0.017218153 | 0.05522743 | 0.047138643 | 0.0814659 | 0.029292 |
| | | *H. sapiens* | *O. anatinus* | *N. chrysocephalum* | *C. porosus* | *C. mydas* | *G. japonicus* | *X. laevis* | *C. milli* |
| **TRAM-TIR** | **Chain B** | | | | | | | | |
| 86 | **A** | 0.061606008 | 0.153759975 | 0.408752851 | 0.215742661 | 0.24865775 | 0.250538455 | 0.3374074 | 0.060994 |
| 87 | **E** | 0.040801468 | 0.382669543 | 0.228637503 | 0.635159367 | 0.40376602 | 0.446224981 | 0.4150428 | 0.037538 |
| 88 | **D** | 0.002893645 | 0.11680307 | 0.281819845 | 0.117765837 | 0.23901556 | 0.075239124 | 0.3451453 | 0.002633 |
| 89 | **D** | 0.172794831 | 0.076095904 | 0.01438122 | 0.042346894 | 0.05395698 | 0.135306708 | 0.2839561 | 0.073681 |
| 116 | **P** | 0.05786548 | 0.338007116 | 0.495613355 | 0.427842124 | 0.28145436 | 0.552244753 | 0.310443 | 0.611771 |
| 117 | **C** | 0.317941098 | 0.579935828 | 0.528464174 | 0.244738697 | 0.32331318 | 0.829997976 | 0.2934107 | 0.76797 |
| 155 | **T** | 0.137402792 | 0.066069031 | 0.055114887 | 0.260483654 | 0.3263984 | 0.140496474 | 0.0421396 | 0.048714 |
| 156 | **S** | 0.572798152 | 0.257495935 | 0.225816061 | 0.531631035 | 0.36592328 | 0.342847776 | 0.3388098 | 0.491543 |
| 167 | **Y** | 0.198116192 | 0.581824885 | 0.06974587 | 0.141088344 | 0.14597639 | 0.146414536 | 0.2605028 | 0.170288 |
| | | *H. sapiens* | *O. anatinus* | *N. chrysocephalum* | *C. porosus* | *C. mydas* | *G. japonicus* | *X. laevis* | *C. milli* |
| **TRIF-TIR** | **Chain C** | | | | | | | | |
| 431 | F | 0.006130579 | 0.000610642 | 0.071973222 | 0.10483366 | 0.1615378 | 0.003100644 | 0.010878 | 0.070015 |
| 440 | S | 0.045661454 | 0.019710541 | 0.004007822 | 0.005244396 | 0.03114083 | 0.001640709 | 0.0518157 | 0.096554 |
| 441 | C | 0.042785443 | 0.023019236 | 0.017149552 | 0.028756832 | 0.01193592 | 0.01654381 | 0.0303221 | 0.019564 |
| 442 | L | 0.11447799 | 0.125205877 | 0.076777481 | 0.134908946 | 0.10299079 | 0.096041277 | 0.1452111 | 0.132049 |
| 443 | Q | 0.032032368 | 0.019952403 | 0.045026998 | 0.01714887 | 0.03776789 | 0.025517827 | 0.0143117 | 0.029171 |
| 518 | Q | 0.550397565 | 0.336690165 | 0.329978637 | 0.968071595 | 0.38594344 | 0.340853117 | 0.1730347 | 0.287843 |
| 519 | I | 0.779633926 | 0.134491506 | 0.040853027 | 0.913541735 | 0.37334953 | 0.39781138 | 0.8884094 | 0.464085 |
| 522 | R | 0.216002877 | 0.094666298 | 0 | 1 | 0.18838453 | 0.058121687 | 0.7444021 | 1 |
| 523 | K | 0.696216208 | 0.464560048 | 0 | 0.933267723 | 0.76742683 | 0.471298536 | 0.5368943 | 0.229183 |

**Figure S8:** The centrality measures highlighting the persistence nature of important residues across organisms for trimeric complexes

| residue no. | residue | *H. sapiens* | *O. anatinus* | *N. chrysocephalum* | *C. porosus* | *C. mydas* | *G. japonicus* | *X. laevis* | *C. milli* |
|---|---|---|---|---|---|---|---|---|---|
| **TRAM-TIR** | **Chain A** | | | | | | | | |
| 86 | **A** | 0.229401465 | 0.339995732 | 0.353241086 | 0.238239605 | 0.439490044 | 0.36588262 | 0.240047 | 0.341101 |
| 87 | **E** | 0.261330113 | 0.22078003 | 0.328783854 | 0.254903913 | 0.436696369 | 0.382816765 | 0.211433 | 0.611664 |
| 88 | **D** | 0.191198589 | 0.386936759 | 0.442673801 | 0.456171494 | 0.757530475 | 0.5134222 | 0.521475 | 0.74025 |
| 89 | **D** | 0.651427847 | 0.268132241 | 0.436920162 | 0.359345273 | 0.611950468 | 0.562410888 | 0.546321 | 0.805437 |
| 116 | **P** | 0.568790434 | 0.304248206 | 0.371904187 | 0.302684421 | 0.247549696 | 0.31793085 | 0.451718 | 0.196132 |
| 117 | **C** | 0.281967645 | 0.190592785 | 0.105945267 | 0.355098878 | 0.287987975 | 0.165479786 | 0.154982 | 0.254048 |
| 155 | **T** | 0.00425682 | 0.008988993 | 0.001916858 | 0.031779637 | 0.038464518 | 0.013316884 | 0.020057 | 0.007727 |
| 156 | **S** | 0.075654232 | 0.05217143 | 0.040128986 | 0.179094202 | 0.060071468 | 0.132700793 | 0.130493 | 0.070419 |
| 167 | **Y** | 0.140037153 | 0.020928361 | 0.03482438 | 0.008870926 | 0.018106776 | 0.041514478 | 0.065768 | 0.016425 |
| | | *H. sapiens* | *O. anatinus* | *N. chrysocephalum* | *C. porosus* | *C. mydas* | *G. japonicus* | *X. laevis* | *C. milli* |
| **TRAM-TIR** | **Chain B** | | | | | | | | |
| 86 | **A** | 0.143385433 | 0.290042372 | 0.141310935 | 0.231862455 | 0.28501237 | 0.410273174 | 0.378934 | 0.056576 |
| 87 | **E** | 0.163901192 | 0.248670829 | 0.173897441 | 0.194202576 | 0.219614417 | 0.339238474 | 0.47518 | 0.05453 |
| 88 | **D** | 0.010710824 | 0.123173069 | 0.125891566 | 0.109456215 | 0.35563534 | 0.205696201 | 0.292284 | 0.003438 |
| 89 | **D** | 0.080161145 | 0.056151437 | 0.043457079 | 0.040466856 | 0.196774457 | 0.033758953 | 0.107572 | 0.073379 |
| 116 | **P** | 0.257491739 | 0.232969576 | 0.239542173 | 0.387655272 | 0.218583513 | 0.517506762 | 0.161096 | 0.525177 |
| 117 | **C** | 0.35714727 | 0.229302702 | 0.235577603 | 0.225103 | 0.520511213 | 0.523944026 | 0.330444 | 0.555003 |
| 155 | **T** | 0.651526783 | 0.034908056 | 0.327365059 | 0.259864864 | 0.316670612 | 0.538039213 | 0.131429 | 0.284766 |
| 156 | **S** | 0.631870713 | 0.237600847 | 0.318213899 | 0.340630265 | 0.475629983 | 0.588100119 | 0.75608 | 0.558891 |
| 167 | **Y** | 0.243752387 | 0.02994271 | 0.033991679 | 0.096313804 | 0.61249913 | 0.054747386 | 0.142268 | 0.54666 |
| | | *H. sapiens* | *O. anatinus* | *N. chrysocephalum* | *C. porosus* | *C. mydas* | *G. japonicus* | *X. laevis* | *C. milli* |
| **TRIF-TIR** | **Chain C** | | | | | | | | |
| 431 | **F** | 0.002334247 | 0.001239846 | 0.004901587 | 0.041087367 | 0.037734925 | 0.013695614 | 0.001518 | 0.232871 |
| 440 | **S** | 0.065495743 | 0.059698768 | 0.06046083 | 0.00665479 | 0.011434527 | 0.011444877 | 0.038372 | 0.101679 |
| 441 | **C** | 0.093174227 | 0.036057661 | 0.065493035 | 0.015175153 | 0.041298889 | 0.024128851 | 0.056956 | 0.03254 |
| 442 | **L** | 0.13782499 | 0.086030421 | 0.132029509 | 0.081508991 | 0.216672504 | 0.095261223 | 0.09256 | 0.269573 |
| 443 | **Q** | 0.115944006 | 0.024014351 | 0.085435715 | 0.041635267 | 0.046457312 | 0.0865137 | 0.069397 | 0.07186 |
| 518 | **Q** | 0.025290671 | 0.104946364 | 0.381628796 | 0.00616941 | 0.070018578 | 0.171247746 | 0.191667 | 0.211854 |
| 519 | **I** | 0.491982165 | 0.386647066 | 0.447555849 | 0.947422573 | 0.634719267 | 0.518024833 | 0.682902 | 0.564364 |
| 522 | **R** | 0.439834216 | 0.254107773 | 0 | 0.528111455 | 0.459414199 | 0.300274528 | 0.225816 | 0.215377 |
| 523 | **K** | 0.437366554 | 0.220507086 | 0 | 0.468925664 | 0.338719123 | 0.51570929 | 0.303539 | 0.501638 |
| | | *H. sapiens* | *O. anatinus* | *N. chrysocephalum* | *C. porosus* | *C. mydas* | *G. japonicus* | *X. laevis* | *C. milli* |
| **TRIF-TIR** | **Chain D** | | | | | | | | |
| 431 | **F** | 0.052288974 | 0.00831264 | 0.017122931 | 0.002008544 | 0.00578386 | 0.000747623 | 0.004582 | 0.00388 |
| 440 | **S** | 0.165574531 | 0.065983209 | 0.037634831 | 0.005562109 | 0.009858938 | 0.009460082 | 0.007184 | 0.0529 |
| 441 | **C** | 0.113607559 | 0.055420795 | 0.027401144 | 0.003447177 | 0.021751746 | 0.015558352 | 0.051433 | 0.019168 |
| 442 | **L** | 0.205205673 | 0.076192106 | 0.062982055 | 0.043318802 | 0.057411798 | 0.088854364 | 0.098415 | 0.091749 |
| 443 | **Q** | 0.057410092 | 0.027444521 | 0.029769256 | 0.015297863 | 0.058394245 | 0.027191565 | 0.018531 | 0.021133 |
| 518 | **Q** | 0.524630298 | 0.332335497 | 0.224936572 | 0.013669297 | 1 | 0.409780806 | 0.305949 | 1 |
| 519 | **I** | 0.268415218 | 0.964768649 | 0.237112587 | 0.189653725 | 0.697403293 | 0.206641466 | 0.330876 | 0.502255 |
| 522 | **R** | 0.435038084 | 0.314044849 | 0 | 0.520658133 | 0.344529368 | 0.210301525 | 0.410164 | 0.694017 |
| 523 | **K** | 0.706760556 | 0.570358395 | 0 | 0.429303809 | 0.561496626 | 0.189082858 | 0.368667 | 0.520156 |

**Figure S9:** The centrality measures highlighting the persistence nature of important residues across organisms for tetrameric complexes

**Modelling attempts for TRAM dimer TRIF monomer complexes trimer model**

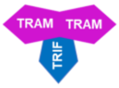

**Model 1**

***1. Protein complex model***
Alpha fold model docked as per known interacting residues form literature using HADDOCK (**guided**)

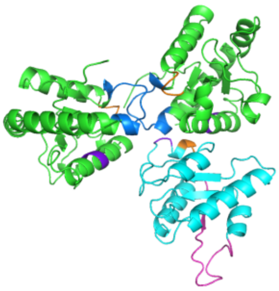

**Model 2**

***2. Protein complex model***
TRAM dimer complex final frame structure from MD (200ns) data followed by **blind** docking using HDOCK

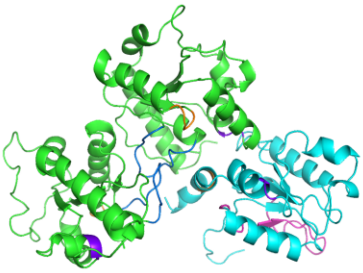

**Model 3**

***3. Protein complex model***
Structure modelling using sequence followed by **blind** docking using HDOCK

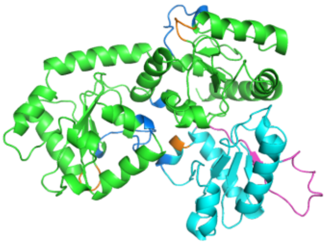

**Model 4**

***4. Protein complex model***
Alphafold multimer modelling (**blind**)

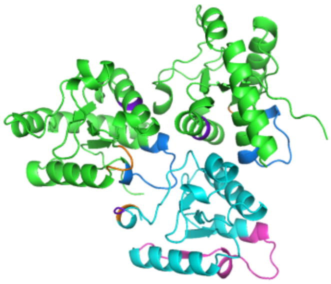

**Model 5**

***5. Protein complex model***
TRAM dimer complex final frame structure from MD (200ns) data followed by **guided** docking using HADDOCK

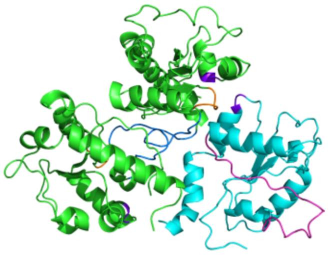

**Model 6**

***6. Protein complex model***
TRAM dimer complex final frame structure from MD (200ns) data followed by **guided** docking using HDOCK

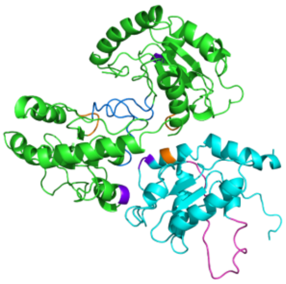

| | MODEL 1 | MODEL 2 | MODEL 3 | MODEL 4 | MODEL 5 | MODEL 6 |
|---|---|---|---|---|---|---|
| Hydrogen Bond Energy (kJ/mol) | -8.63 | -30.84 | -31.22 | -63.69 | 0.00 | -19.73 |
| Electrostatic Energy (kJ/mol) | -51.19 | 2.78 | -15.52 | -0.74 | -20.52 | -45.56 |
| Van der Waals Energy (kJ/mol) | -189.26 | -262.28 | -243.53 | -310.59 | -108.93 | -143.93 |
| **Total Stabilizing Energy** (kJ/mol) | **-249.08** | **-290.35** | **-290.27** | **-375.02** | **-129.45** | **-209.22** |
| Number of interface residues | 93 | 98 | 110 | 111 | 73 | 101 |
| **Normalized Energy per residue (kJ/mol)** | **-2.68** | **-2.96** | **-2.64** | **-3.38** | **-1.77** | **-2.07** |
| No. of Short Contacts | 7 | 13 | 28 | 12 | 2 | 47 |
| No. of Hydrophobic Interactions | 3 | 2 | 0 | 4 | 2 | 1 |
| No. of van der Waals Pairs | 7821 | 9591 | 9730 | 11805 | 2928 | 8864 |
| No. of Salt Bridges | 2 | 1 | 1 | 1 | 1 | 3 |
| No. of Potential Favourable Electrostatic Interactions | 4 | 4 | 6 | 3 | 6 | 10 |
| No. of Potential Unfavourable Electrostatic Interactions | 1 | 9 | 4 | 4 | 3 | 3 |

**Figure S10:** The structure of the various trimeric complex models and its energies
